# Coordinated control of neuronal differentiation and wiring by a sustained code of transcription factors

**DOI:** 10.1101/2022.05.01.490216

**Authors:** Mehmet Neset Özel, Claudia Skok Gibbs, Isabel Holguera, Mennah Soliman, Richard Bonneau, Claude Desplan

## Abstract

The enormous diversity of cell types in nervous systems presents a challenge in identifying the genetic mechanisms that encode it. Here, we report that nearly 200 distinct neurons in the *Drosophila* visual system can each be defined by unique combinations of ∼10 transcription factors that are continuously expressed by them. We show that targeted modifications of this ‘selector’ code induce predictable conversions of cell fates between neurons *in vivo*. These conversions appear morphologically and transcriptionally complete, arguing for a conserved gene regulatory program that jointly instructs both the type-specific development and the terminal features of neurons. Cis-regulatory sequence analysis of open chromatin links one of these selectors to an upstream patterning gene in stem cells that specifies neuronal fates. Experimentally validated network models show that selectors interact with ecdysone signaling to regulate downstream effectors controlling brain wiring. Our results provide a generalizable framework of how specific fates are initiated and maintained in postmitotic neurons.

## Introduction

Neurons are by far the most diverse of all cell types in animals. A recent single-cell RNA sequencing (scRNA-seq) survey of the mouse cortex and hippocampus, for instance, has identified 364 distinct neuronal types^1^ and this is likely a gross underestimation^2^. Understanding the molecular mechanisms that generate and maintain this vast diversity is a central goal of neurobiology. The *Drosophila* brain provides a tractable system to approach this challenge due to its manageable size (∼100,000 neurons), genetically hardwired (i.e., experience-independent) development, and powerful genetic tools. The fly brain can be divided into two main regions: The central brain constitutes about a quarter of the entire brain and contains more than 5,000 morphologically and connectomically unique neuron types^3^. This diversity remains largely unresolvable by scRNA-seq^4^, since many of these types exist in only two cells per brain. The optic lobes contain the majority of neurons in the brain at significantly lower diversity (∼200 types). Each of their neuropils: lamina, medulla, lobula and lobula plate (**Fig. 1A**) is divided into ∼800 columns, corresponding to the same number of ommatidia (unit eyes) in the retina. Because of this retinotopic organization with multiple repeats of the same circuits, most neurons are present at a much higher number of cells per type. We recently completed a large scRNA-seq atlas of the *Drosophila* optic lobes, resolving around 200 cell types that we consistently tracked across 6 timepoints from early pupal stages to adult^5^. Importantly, almost all annotated clusters in this atlas correspond to a distinct neuronal type with unique morphology^6^. This strongly suggests that the vast majority of our clusters represent biologically homogeneous groups, giving us access to the cell-type specific transcriptome of every optic lobe neuron throughout its development.

**Figure 1:**
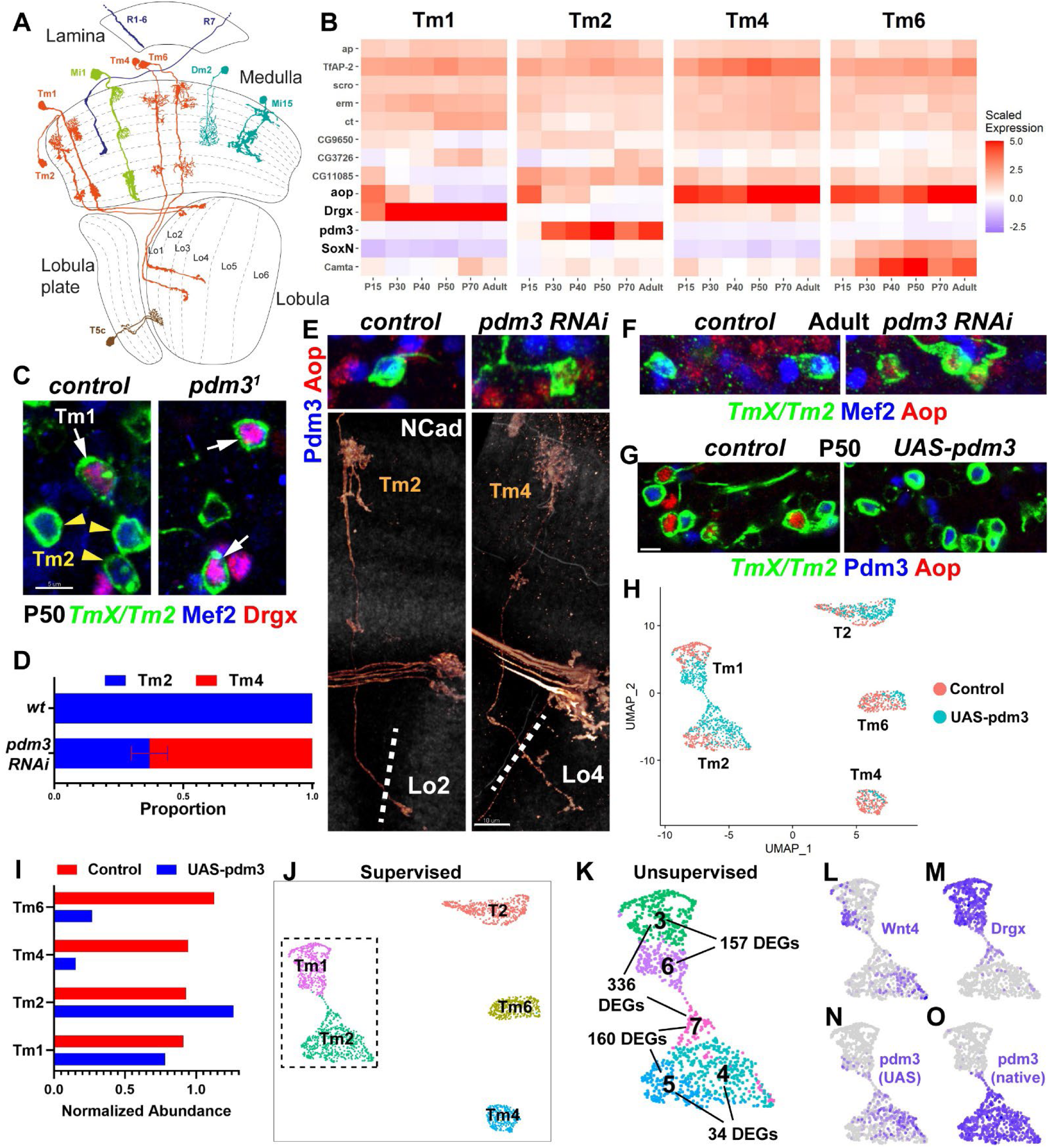
Selector TF *pdm3* instructs complete switches of neuronal fates. **A,** Ventral view of the *D. melanogaster* optic lobe in cross-section, with drawings of select cell types. Dashed lines: boundaries between layers. Partially adapted from^6^. **B,** Developmental (scaled) expression patterns of all genes that are candidate selectors in either of the displayed cell types (Fig. S1A), based on^5^. Note that scaling was performed over the entire atlas at each stage, thus the values are comparable between the cell types, but not stages. Colormap max. value was capped at 5 to preserve dynamic range. **C,** FRT40A (control) and *pdm3*^1^ MARCM clones labeled with *TmX/Tm2-Gal4* and CD4tdGFP in P50 brains (maximum projection), with co-stainings of anti-Mef2 (blue) and anti-Drgx (red). Only Mef2^+^ neurons remaining in mutant clones are the Drgx^+^ Tm1 (n= 8 brains for control and 4 brains for mutant). **D-F**, *TmX/Tm2-Gal4* driving *pdm3* RNAi and UAS-CD4tdGFP (flip-out). **D,** Quantification of E-F (combined). n= 54 neurons in 6 brains (control), 96 neurons in 8 brains (RNAi). p<0.0001 for change in Tm4 proportion. Error bar denotes SEM. **E,** 3D reconstructions of GFP expression (bottom) or max projections (top) for the same representative adult neurons in each condition, with co-stainings of anti-NCad (marking neuropils, bottom), anti-Pdm3 (blue) and anti-Aop (red) (top). The dashed lines indicate the border of the lobula neuropil based on NCad staining (see Fig. S2D), which is displayed weakly to avoid obstructing the details of morphology. **F,** Same as (E, top) with co-staining of anti-Mef2 instead of Pdm3. **G,** *TmX/Tm2-Gal4* driving *UAS-pdm3* and UAS- CD4tdGFP (flip-out). Max. projections of cell bodies in P50 medulla cortex, with co-stainings of anti-Pdm3 (blue) and anti-Aop (red). Aop^+^ neurons are almost completely eliminated with *UAS- pdm3* (n=118 neurons in 12 brains). Scale bars: 5 μm (C, E-top, F, G) and 10 μm (E-bottom). **H-O,** scRNA-seq of FACSed neurons, same experiment as (G) with nuclear GFP labeling. UMAP visualizations of the displayed cell types were calculated using their top 6 principal components. Cells are colored according to library (condition) of origin (**H**), neural network^5^ classifications (**J**), unsupervised clustering (**K**, inset only, see also Fig. S2H, DEGs: differentially expressed genes, see Table S2) and the log-normalized expression values of indicated genes (**L-O**). **I**, Numbers of Tm neurons (according to neural network classifications) in each library, divided by the number of T2 neurons in the same library. Note that in (**L**), Wnt4 can be seen in both cluster 4 (converted Tm4/6) and cluster 6 (Tm1 with ectopic *pdm3*). Expression in cluster 6 may indicate that co-expression of Drgx and Pdm3 creates a novel regulatory state, since neither Tm1 nor Tm2 normally express Wnt4.

The identity of optic lobe neurons is specified deterministically by their progenitors during neurogenesis that occurs from late larval stages (L3) until about 20% of pupal development (P20)^7^. Neurons from the largest neuropil of the optic lobe, the medulla, are produced from a neuroepithelium called the outer proliferation center (OPC), which is progressively converted into neural stem cells (neuroblasts) that asymmetrically divide multiple times, each time self-renewing and producing a Ganglion Mother Cell (GMC) that divides once to generate two different neurons^8^. Intersection of three patterning mechanisms diversify the OPC progeny: Compartmentalization of the neuroepithelium into at least 8 spatial regions by transcription factors (TFs) and signaling molecules^9^, sequential expression of at least 11 temporal TFs (tTFs) in neuroblasts^10^, and differential Notch signaling between the GMC daughters^11, 12^. Similar temporal and spatial patterning mechanisms are also utilized in other parts of the fly brain, as well as mammalian neural stem cells to generate diversity (reviewed in^13^). Although we understand the mechanisms that initially specify neuronal type identity, how these cell fate decisions are then implemented and maintained in postmitotic neurons remains much less clear. Faithful retention of cell identity is critical for every neuron throughout its life, so that it can acquire its unique morphology and connectivity during development, and express its correct synaptic and electrical machinery in the functioning brain. Specific genetic programs must exist to coordinate these since most spatial and tTFs are not maintained in neurons^10^. These genetic programs also need to interact with extrinsic signals^14^ to ensure that correct downstream “effector” genes are activated at the correct times.

Much of our knowledge about neuronal identity control originates from *C. elegans*. The terminal selector hypothesis^15^ posits that most, if not all, *type-specific* gene expression in neurons is controlled by combinations of TFs that are *continuously maintained* in each neuron throughout its life. These terminal selectors control both the developmental features such as synaptic connectivity^16, 17^ and the functional features such as neurotransmitter identity^18, 19^, but they are not required for the pan-neuronal gene expression programs^20, 21^. An important implication of this model is that individual terminal selectors do not specialize in distinct phenotypic features of a neuron. Effector genes are controlled by different *permutations* of available selectors in a given neuron, which implies that every effector gene might not be regulated by all terminal selectors. Although a few TFs that could function as terminal selectors have been identified in mammalian neurons^22–24^, it remains unclear how generally applicable this regulatory logic is beyond the relatively simple nervous system of worms. For instance, some TFs were postulated to regulate the morphological, but not the terminal, features of *Drosophila* leg motoneurons^25^. Moreover, the ultimate test of this model, *i.e.* the *predictive* and *complete* transformation of one neuronal type into another through targeted modification of its selector code, has been difficult to assess, even in *C. elegans*^26^.

In this study, we took advantage of our developmental scRNA-seq atlases to identify the potential terminal selector genes, which we refer here as “selector TFs” (see Discussion), in all optic lobe neurons. We found unique combinations of, on average, 10 TFs differentially expressed and stably maintained in each neuron. Through genetic gain- and loss-of-function experiments in postmitotic medulla neurons, we show that modifications of these selector TF codes are sufficient to induce predictable switches of identity between 6 different cell types. These TFs jointly control both developmental (*e.g.* morphology) and functional (*e.g.* ion channels, neurotransmitters) features of neurons. scRNA-seq of specific perturbed neurons indicate that type conversions are essentially complete, not only morphologically but also transcriptionally. We also show that the selector TF Drgx in a specific medulla neuron (Tm1) is regulated by the tTF Klumpfuss that instructs its fate in progenitors, establishing a link between the regulatory programs that specify and maintain neuronal fates. Finally, we combined our transcriptomic datasets with chromatin accessibility datasets^27^ to understand the mechanisms that control brain wiring downstream of selector TFs, by building predictive computational models of gene regulatory networks. Experimental validations of these networks revealed how selector TFs interact with ecdysone-responsive TFs to activate a large and specific repertoire of cell-surface proteins and other effectors in each neuron at the onset of synapse formation. Together, our results provide a unified framework of how specific fates are initiated and maintained in postmitotic neurons and open new avenues to understand synaptic specificity through gene regulatory networks.

## Results

### Selector transcription factors of optic lobe neurons

To investigate if a sustained code of TFs maintains the identity of each neuron throughout development, we sought to identify the combinations of candidate selector TFs expressed in each of the 174 neuronal clusters in our scRNA-seq atlas^5^. We determined the sets of TFs continuously expressed in each cluster throughout all six stages (P15-Adult) that were previously profiled, excluding those expressed in all clusters (pan-neuronal or ubiquitous genes, see Methods). We found on average unique combinations of 10 such genes per cluster, representing 95 TFs in total (**Fig. S1A**, **Table S1**). This sustained TF code could encode neuronal identity on its own. Homeobox genes were enriched in this list (**Fig. S1B**) compared to other TF classes, but unlike in the *C. elegans* nervous system^28^, homeobox genes alone were not sufficient to uniquely define every neuron in the optic lobe. It is important to note that some of these TFs were expressed continuously in some neurons but only transiently in others; they were only considered as part of the selector code in the former case.

The most compelling prediction of the selector TF hypothesis is that, if continuously maintained TFs are primarily responsible for cell type-specific neuronal differentiation, it should be possible to engineer predictable switches of identity between different neurons by modifying these TFs alone. Remarkably, all the genes that have so far been reported to interfere with neuronal type identity in the *Drosophila* optic lobe: *bsh* in Mi1, L4 and L5 neurons^29^, *hth* in Mi1 neurons, *drifter* (*vvl*) in Tm27Y neurons^30^, *hth* and *Lim1* in Lawf2 neurons^31^, *erm* in L3 neurons^32^, and *SoxN* and *Sox102F* in T4/5 neurons^33^, are defined in our atlas as selector TFs for each of these neurons (**Fig. S1A**, **Table S1**). However, these studies have generally reported disruptions rather than switches of morphological identity: Although loss of *hth*/*bsh* in Mi1 generates “Tm1-like” neurons^30^, and loss of *bsh* in L4 leads to “L3-like” neurons^29^, the perturbed neurons are clearly distinguishable from wild-type Tm1 and L3. In hindsight of the selector TF code we propose here, these incomplete conversions would have been expected: Mi1 and Tm1 can also be distinguished by differential expression of other selector TFs, *Drgx* and *TfAP-2* in Tm1, while L3 and L4 are also distinguished by *erm* (L3) and *tj* (L4) (**Fig. S1A**). However, it remains challenging to simultaneously perturb more than one or two genes at once using classical genetic methods. In order to provide definitive evidence for the sufficiency of selector TFs in determining neuronal type identity, we thereby looked for groups of closely related neurons whose selector codes differ only by one or two genes, where complete conversions from one cell-type to another may be feasible.

We focused on Transmedullary (Tm) neurons 1, 2 and 4, which are members of the OFF-pathway critical for detection of wide-field motion: They relay information from the L2 and L3 neurons in the medulla to the motion-selective T5 neurons in the lobula^34, 35^. Although these Tms share similar overall morphology, adult neurons are readily distinguishable from one another by their distinct dendritic shapes as well as the different target layers of their axons in the lobula (**Fig. 1A**). At P15, shortly after their terminal division, these three neurons, along with an unidentified cluster (#62), have nearly indistinguishable transcriptomes^5^, suggesting a very close developmental relationship. We annotated cluster 62 as Tm6 neurons, based on its expression of the unique combination of *aop, SoxN* and *Wnt10* (**Fig. S2A-C**). Analysis of candidate selector TF expression in these clusters across development (**Fig. 1B**) revealed that the 4 neurons indeed share a very similar code: *ap*, *TfAP-2*, *scro*, *erm* and *ct* are continuously expressed in all four clusters. *CG9650*, *CG3726, CG11085* and *aop* could also be found in all 4 Tm neurons at some point during development, though they are only transiently expressed in some of them (*e.g.,* Tm4 and Tm6 continuously express *aop*, but these transcripts are also present in Tm1 and Tm2 until P40). *Camta* is expressed at much higher levels in Tm6 but is also detected in the others. While there are no candidate selectors exclusive to Tm4, among these 4 neurons, *Drgx* is specific to Tm1, *pdm3* to Tm2 and *SoxN* to Tm6. Therefore, these TFs that are each continuously and specifically expressed in one of these types are strong candidates to differentially specify their fates.

### *Pdm3* instructs transcriptionally complete switches of neuronal fate

Pdm3 is important for the development of several neurons in the *Drosophila* brain^36–38^, but it has not been studied in the visual system. As Pdm3 is the only TF that continuously distinguishes Tm2 from Tm4 (**Fig. 1B**), the selector TF code predicts that its loss should reprogram Tm2 neurons to Tm4 fate. *R71F05-Gal4* is expressed in all 4 Tm neurons (1/2/4/6) until P50; however, it is only maintained in Tm2 in adults (**Fig. S2D**). We henceforth refer to it as *TmX/Tm2-Gal4*. Using this driver, we generated MARCM^39^ clones of a *pdm3* null allele^37^. No mutant Tm neurons were recovered in adult brains, suggesting that the Gal4 was turned off, and thus that Tm2 were not specified properly (**Fig. S2E**). *Mef2* is an effector (downstream) TF normally expressed specifically in both Tm1 and Tm2 after P40 (**Fig. 1C**, **Fig. S5F**). At P50, we observed that the only remaining Mef2^+^ Tm neurons in *pdm3* mutant clones were Tm1 that expressed Drgx (**Fig. 1C**), indicating that Tm2 were either lost or converted to another fate. Interestingly, upon RNAi knock-down of *pdm3* using *TmX/Tm2-Gal4*, 65% of Tm2s that kept the expression of the driver in adult brains were converted to neurons with Tm4 morphology, characterized by wider dendritic arbors that are symmetrical around the main fiber of the neuron as well as axons targeting to the deeper lobula layer 4 (**Fig. 1D-E**, compare to **1A**). It is likely that the knock-down retains low levels of Pdm3 in Tm2 that are sufficient to maintain expression of *TmX/Tm2-Gal4* in adults but are insufficient for instructing the Tm2 fate. These Tm4-looking neurons did not express Mef2 and instead expressed the putative Tm4 selector TF Aop (**Fig. 1E-F**). This could argue that Aop^+^ Tm4 is a “ground state” among these neurons that is overridden by Pdm3 in Tm2, which we validate further in the following sections.

We then asked whether ectopic expression of *pdm3* in Tm4 and Tm6 could be sufficient to convert them to Tm2 fate. We used *TmX/Tm2-Gal4* to express *UAS-pdm3.short*^36^ and found that Aop^+^ neurons (Tm4 and Tm6) were almost completely eliminated at P50 (**Fig. 1G**), suggesting that they had been lost or converted. Furthermore, performing this experiment with a Tm4/6-specific driver led to the nearly complete loss of Gal4 expression in adults (**Fig. S3C,** shown is the only case of a labeled neuron). To address the completeness of these conversions at P50, when the neurons have not fully acquired their adult morphology but display the greatest transcriptomic diversity^5^, we analyzed their gene expression. We labeled all 4 Tm neurons (*TmX/Tm2-Gal4*) with nuclear GFP, crossed to either *UAS-pdm3* or *yw* (control), isolated the labeled cells using FACS, and performed single-cell RNA sequencing. We acquired 6693 cells passing QC filters and annotated them (**Fig. S2F**) using a neural-network classifier trained on our reference atlas of the optic lobe^5^. As the driver weakly labels several cell-types other than Tm neurons (see Methods), we only retained the 1779 cells (1056 control and 723 *UAS-pdm3*) classified as Tm1/2/4/6, in addition to 563 classified as T2 that are also strongly labeled by this driver (**Fig. 1H,J**). T2 neurons, like Tm2, natively express *pdm3*, and thus they should not be affected by this perturbation and serve as an internal control. We calculated the relative abundance of each Tm neuron, normalized by the number of T2 in the corresponding library. As expected, we observed a dramatic depletion of Tm4 and Tm6 in the UAS-pdm3 library compared to control and an increase in the number of Tm2s (**Fig. 1I**), validating that ectopic *pdm3* converts Tm4 and Tm6 to Tm2.

We noted that the increased number of Tm2s upon *pdm3* overexpression was not sufficient to fully account for the lost Tm4s and Tm6s. Some optic lobe neurons are known to be generated in excess, followed by widespread apoptosis in the first half of pupal development^40^ as neurons that fail to match with their correct partners are eliminated^41^. Thus, many of the supernumerary Tm2s may be undergoing programmed cell death. We stained these brains using an antibody against cleaved Dcp-1, an activated caspase that marks apoptotic cells shortly before they die^42^. In control brains at P25, 3.1% of all cells labeled by GFP (*TmX/Tm2-Gal4*) in the medulla cortex were apoptotic (n=73/2313 cells across 5 brains), but none of these were Tm2s (0/561, based on GFP/Pdm3 colocalization). In brains overexpressing *pdm3*, 6.1% of all GFP+ cells (n=163/2640 cells across 5 brains, p=0.006) as well as 5.9% of GFP+/Pdm3+ cells (128/2157, p<0.0001) were undergoing apoptosis (**Fig. S2G**). Together, these results suggest that when excess Tm2 neurons are produced through conversions from Tm4 and Tm6, this is compensated by increased cell death. This mechanism potentially helps ensure that only one Tm neuron of each type is present per column in wild-type brains.

To distinguish the wild-type Tm2 neurons from those converted from another cell type, we performed unsupervised clustering on the dataset. This revealed heterogeneous populations among the cells classified as Tm1 and Tm2 (**Fig. S2H**, **Fig. 1K**). Tm2 subclusters 4 and 5 were extremely similar, with only 34 significant differentially expressed genes (DEGs) between them (Methods). Most of those 34 DEGs were subtle differences in levels (**Table S2**), but the strongest marker was *Wnt4*, found in cluster 4 (**Fig. 1L**). We recently showed that *Wnt4* is expressed in ventral Tm4 and Tm6, but not in Tm2^5^, strongly suggesting that cells in cluster 4 were converted neurons that had retained Wnt4 expression from their initial specification as Tm4/6. Nevertheless, these differences between the converted and ‘original’ Tm2s were minimal, as compared to more than 700 DEGs normally observed between wild-type Tm2 and Tm4 at this stage (**Table S2**). We therefore conclude that conversion from Tm4/6 to Tm2 induced by ectopic *pdm3* appears complete.

The third subgroup of the cells classified as Tm2, cluster 7, consisted entirely of cells from the UAS-pdm3 library and surprisingly also expressed the Tm1 selector *Drgx* (**Fig. 1M)**, suggesting that these were originally Tm1s converted to a Tm2-like state. Despite being classified as Tm2s, they were significantly different from cluster 5 (wt Tm2) with 160 DEGs (**Fig. 1K**). The cells classified as Tm1 were clustered into two groups: cluster 6, made entirely of cells from the UAS-pdm3 library, and was significantly different from cluster 3 that consisted essentially of wt Tm1s (**Fig. 1H,K**). Thus, both clusters 6 and 7 contained Tm1s with ectopic *pdm3*. UMAP visualization showed a thin stripe of cells bridging the Tm1-like (cluster 6) and Tm2-like (cluster 7) states. We investigated the dynamics of this transition. Interestingly, while both clusters 6 and 7 displayed reads coming from the *UAS-pdm3* construct as expected (**Fig. 1N**), cluster 7 also expressed *pdm3* from the native locus (**Fig. 1O**). We thereby conclude that the amount of protein produced from the UAS construct is insufficient for conversion into Tm2, but instead Pdm3 must autoactivate above a certain threshold. Once this threshold is reached, Pdm3 quickly drives Tm1 and Tm4/6 (that also never normally express *pdm3*) to a Tm2-like state; however, this conversion remains incomplete in Tm1 (cluster 7) since *Drgx* remains expressed **(Fig. 1M)**. This positive feedback additionally explains why *pdm3* overexpression was able to turn off expression of a Tm4/6-specific Gal4 in adults (**Fig. S3C**): once the autoactivation threshold was reached, the conversion was permanent and the Gal4 driven *pdm3* was no longer required. Morphologically, Tm1 neurons overexpressing *pdm3* appeared normal in adults despite co-expressing Drgx and Pdm3, which we validated by immunostainings (**Fig. S2I**). We suspect that these neurons correspond to the cells of cluster 6 in the scRNA-seq experiment, suggesting that the 157 DEGs between clusters 3 (wt Tm1) and 6 are not important for morphology. Tm2-like cluster 7 cells have most likely turned off Tm1-Gal4 expression as it is unlikely that a transcriptomic shift of that extent (336 DEGs at P50) would yield morphologically normal Tm1s.

In summary, *pdm3* is necessary and sufficient to instruct the fate choice between Tm2 and Tm4 neurons, as predicted by the selector TF code. Its loss results in morphological conversion of Tm2 into Tm4, and its ectopic expression can induce essentially complete transcriptomic conversions of Tm4 and Tm6 to Tm2 fate. It is also an upstream repressor of the Tm4/6 selector TF *aop* (**Fig. 1E,G**).

### The temporal TF Klumpfuss regulates the selector TF Drgx in Tm1

Similar to *pdm3* in Tm2, Drgx is the only TF that continuously distinguishes Tm1 from Tm4 (**Fig. 1B**). We ectopically expressed *Drgx* using *R35H01-Gal4*, which is expressed in all 4 Tm neurons until P50, but is only maintained in Tm4 and Tm6 in adults (**Fig. S3A-B, Fig. 2A**, *TmX/Tm4,6-Gal4*). In these adult brains, the proportion of Tm6 remained unchanged, but most Tm4s were morphologically converted into Tm1s, characterized by much narrower dendritic arbors and axons terminating in the first layer of the lobula (**Fig. 2A-B**, see **Fig. 1A**). Importantly, the converted neurons also lost Aop expression and instead expressed Mef2 (Tm1/2 marker) (**Fig. 2C**), indicating that conversion was not limited to morphology. Some of the converted neurons displayed morphological features atypical of Tm1, such as targeting to the Lo2 layer instead of Lo1 (**Fig. S3D**). We suspect that this partial expressivity is due to low Gal4 expression from *TmX/Tm4,6-Gal4.* Moreover, the fact that this driver is expressed at even weaker levels in Tm6 compared to Tm4 (**Fig. S3B**) might explain our failure to affect Tm6 fate. Alternatively, the expression of an additional selector (*SoxN*) in Tm6 may confer resistance to this conversion, although Pdm3 misexpression was able to convert Tm6 into Tm2. Loss of *Drgx* (described below) resulted in conversion of Tm1 neurons into Tm4 (**Fig. 2G-H**). Thus, *Drgx* specifies the Tm1 fate; and it can mediate the conversion of Tm4 into Tm1 as predicted by the selector code. It is also a repressor of *aop*, further suggesting that Tm4 might be the ground state that is overridden by other selectors.

**Figure 2:**
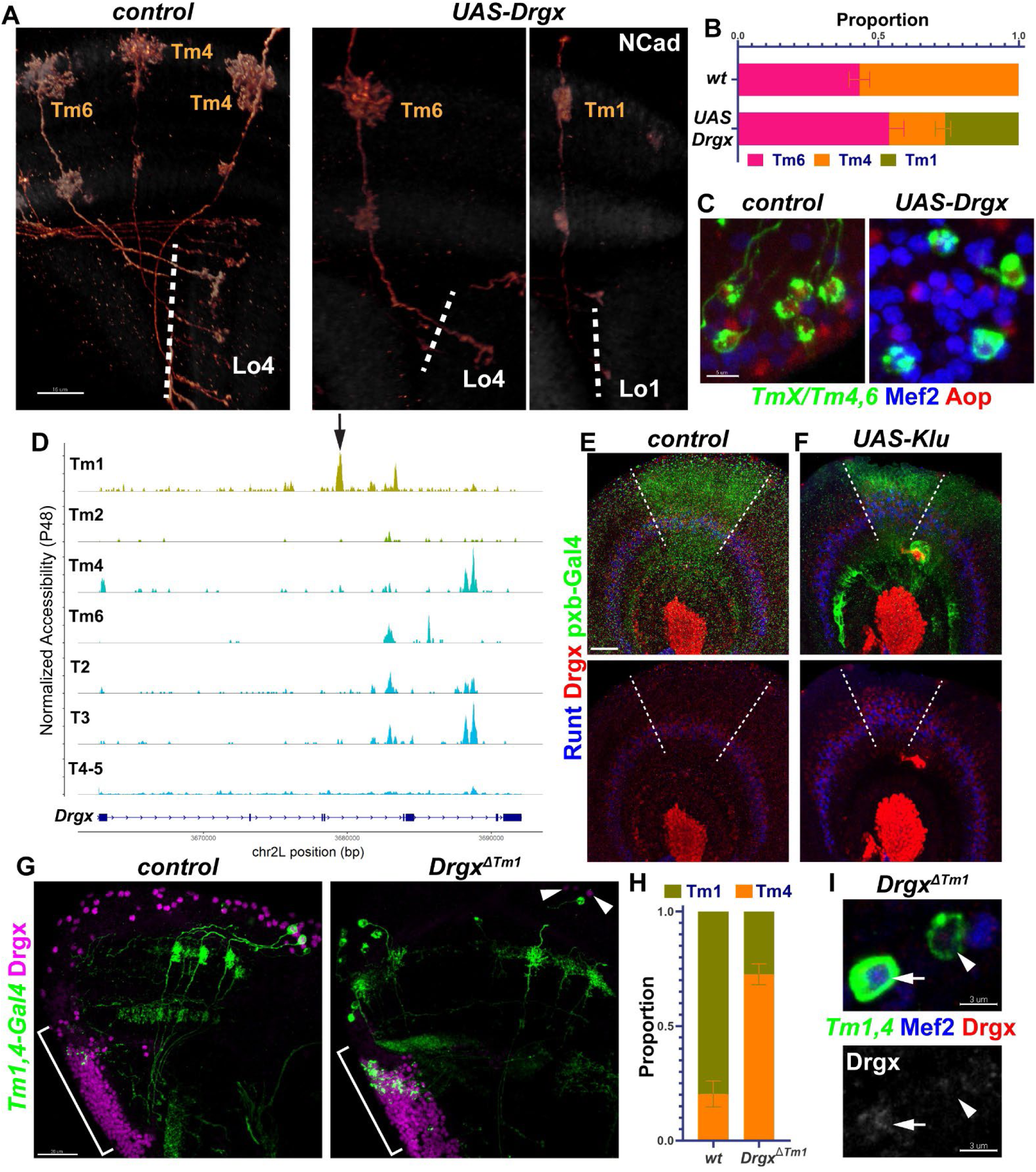
Drgx is a selector TF of Tm1 neurons and is regulated by the tTF Klu. **A-C,** *TmX/Tm4,6-Gal4* driving UAS-Drgx and UAS-CD4tdGFP (flip-out). **A,** 3D reconstructions of GFP expression for representative adult neurons in each condition (see Fig. 1A), with co- staining of anti-NCad (white). Dashed lines mark the border of lobula neuropil. **B,** Quantification of A. n= 92 neurons in 4 brains (control), 45 neurons in 6 brains (Drgx). p=0.0003 for change in Tm1 proportion. **C,** Same as (A) with max. projections of cell bodies with co-stainings of anti- Mef2 (blue) and anti-Aop (red). **D,** Aggregated accessibility tracks of *Drgx* locus for the indicated cell types from the TF-IDF normalized scATAC-seq data at P48^27^. Arrow points to the Tm1- specific enhancer deleted in (G-I). **E-F,** *pxb-Gal4* driving UAS-CD8GFP and UAS-Klu (**F**, n=5 brains) in L3 optic lobes, with co-stainings of anti-Runt (blue) and anti-Drgx (red). Dashed lines mark the borders of driver expression where the Runt/Drgx concentric region is expanded upon Klu overexpression. **G-I,** *Tm1,4-Gal4* driving UAS-CD4tdGFP (flip-out) in the background of heterozygous (control) or homozygous *Drgx^ΔTm1^* allele. **G,** Max projections of adult optic lobes with co-staining of Drgx (magenta). Brackets mark the location of the lobula plate cortex where T2-5 neurons are located. Arrowheads in the mutant point to remaining Drgx^+^ cells in the medulla cortex which are glia (see Fig. S3F). **H,** Quantification of G (see also Fig. S3E) based on morphology of the observed neurons. Note that Tm1 were normally observed more frequently than Tm4 (despite their normally equal stoichiometry) as the driver expression is much lower in Tm4. n = 57 neurons in 6 brains (control) and 181 neurons in 10 brains (*Drgx^ΔTm1^*). p<0.0001 for change in Tm1 or Tm4 proportion. **I,** Same as (G), displaying instead cell bodies with co-stainings of anti-Mef2 (blue) and anti-Drgx (red). Drgx single channel is also shown separately (bottom). Arrow points to a Tm1 neuron that did not convert and maintained Mef2 expression, with low levels of Drgx barely detectable. Arrowhead points to a Tm4 (or a fully converted) neuron. Scale bars: 15 μm (A,E,F), 5 μm (C), 20 μm (G), 3 μm (I). Error bars denote SEM.

Neither *Drgx* nor *pdm3* is expressed in the progenitors (neuroblasts) of Tm neurons^10^, implying that their postmitotic expression in specific neurons is instructed by tTFs in the neuroblasts. To investigate how the expression of selector TFs is controlled, we used a single-cell ATAC-seq (chromatin accessibility) dataset of the developing *Drosophila* brain^27^. We identified the cells belonging to optic lobe neurons at adult, P48 and P24 stages (individual neuronal types were not resolvable at earlier stages), re-clustered and annotated them using our scRNA-seq atlas^5^ as reference (**Fig. S4A-D**, Methods). Analyzing the differentially accessible regions near the *Drgx* locus, we identified a putative enhancer in the 4^th^ intron of the gene that was specifically accessible in Tm1 throughout development (**Fig. S4E**) but was not accessible in the other Tm neurons, or in T2, T3 and T4/5, which also express *Drgx* (**Fig. 2D**). This suggested that this 700bp region is a Tm1-specific regulatory element. We found that the only enriched binding motifs for any of the TFs expressed in the optic lobe (E-value<100, see Methods) within this enhancer, belonged to the tTF Klumpfuss (Klu). Klu is expressed at higher levels in neuroblasts during early temporal windows, when Tm1 is generated, and its overexpression in neuroblasts can expand Runt^+^ neurons^43^ that are likely born in the same temporal window as Tm1 (**Fig. 2E**)^10^. Thus, Klu might also regulate *Drgx* expression. Indeed, *Klu* overexpression using *pxb-Gal4*, which is expressed in the Vsx (central) region of the OPC (**Fig. 2E-F**, dashed lines), resulted in the expansion of Drgx^+^ neurons (i.e. Tm1) in this region, similar to Runt (**Fig. 2F**). We therefore conclude that Klu expression in neuroblasts helps specify Tm1 from an early temporal window by activating the selector TF *Drgx* in their neuronal progeny.

Next, we asked whether *Drgx* expression is regulated by the enhancer element we had identified from scATAC-seq. We engineered a CRISPR deletion of this intronic region (Methods). If this enhancer is required for *Drgx* expression in Tm1, the resulting allele (*Drgx*^ΔTm^^1^) should function as a conditional mutant specifically in these neurons. The homozygous *Drgx*^ΔTm^^1^ flies were fully viable and fertile, as opposed to the presumed-null *Drgx* CRIMIC^44^ that are homozygous lethal. *27b-Gal4*^45^ is expressed in Tm1, and much more weakly in Tm4 (**Fig. S3E**, *Tm1,4-Gal4*) throughout development. In *Drgx*^ΔTm^^1^ mutant adults, Drgx expression in medulla cortex (where all Tm cell bodies are located) was almost completely lost (**Fig. 2G**) but was still normally present in Repo^+^ perineurial glia (**Fig. S3F**) at the surface of the brain (**Fig. 2G**, arrowheads) and in T-neurons originating from the lobula plate (**Fig. 2G**, brackets). The observed ratio of Tm1/Tm4 labeled by *Tm1,4-Gal4* decreased dramatically in the mutant brains (**Fig. 2H**), suggesting that most Tm1s were converted to Tm4 based on morphology. In addition, 69% of the few remaining Tm1s displayed abnormal morphological features such as disrupted dendritic arbors, and/or axons reaching to deeper layers in the lobula (**Fig. S3E**). Close examination of cell bodies revealed that these neurons that maintained Mef2 (Tm1/2 marker) still expressed Drgx at very low levels (**Fig. 2I**, arrow, arrowhead points to a Tm4). Surprisingly, Drgx expression was normal in *Drgx*^ΔTm^^1^ mutants at L3 stage (**Fig. S3G**). These results suggest that there are other, partially redundant enhancers regulating *Drgx* that control its initial activation in nascent neurons, while the robust maintenance of expression in Tm1s requires this specific enhancer. The deletion of this element effectively created a cell-type specific knock-down of *Drgx*, leading to the destabilization of Tm1 fate and stochastic conversions to Tm4.

In summary, Drgx is necessary and sufficient for the specification of Tm1 fate (over Tm4), which is instructed by the temporal expression of Klu in neuroblasts. A differentially accessible enhancer that we identified from scATAC-seq contributes to this specification. Klu could be priming this enhancer in newborn Tm1s, even though it is not maintained later, to ensure sustained expression of *Drgx*. In addition, the *Drgx*^ΔTm^^1^ allele is a powerful proof-of-principle for the generation of targeted, cell-type specific mutants based entirely on high-throughput chromatin accessibility data. With the increasing prevalence of single-cell multiome techniques and CRISPR/Cas9 genome editing, we believe this approach will prove particularly valuable in mammalian systems where targeted disruption of genes in specific cell-types generally requires time-consuming conditional schemes.

### Selector TFs jointly control developmental and functional neuronal type identity

Our results thus far have shown that selector TF combinations determined from scRNA-seq data enable predictive modifications of neuronal type identity that appear to be effectively complete for both morphology and transcriptome. Similar to *Drgx* in Tm1 and *pdm3* in Tm2, *SoxN* is the sole candidate selector distinguishing Tm6 neurons from Tm4 (**Fig. 1B**). In control MARCM clones marked with *TmX/Tm4,6-Gal4*, we observed roughly equal numbers of adult Tm4 and Tm6. In contrast, only Tm4s could be observed in *SoxN* null mutant^46^ clones (**Fig. 3A**). All four Tms are unicolumnar neurons produced by all neuroblasts; therefore, a given column should contain one Tm4 and one Tm6 generated from the same neuroblast^9^. We consistently observed Tm4 and Tm6 neurons occupying the same column in sparse control MARCM neuroblast clones (**Fig. 3B**). However, these clones consisted of two Tm4s in *SoxN* mutants (**Fig. 3B**), indicating that loss of *SoxN* converted Tm6 neurons to Tm4, rather than eliminating them. However, columns with these pairs were rare, suggesting that the extra Tm4s often undergo apoptosis as shown above for Tm2s. Overexpressing *SoxN* using *TmX/Tm4,6- Gal4* did not convert Tm4s to Tm6 (**Fig. S5A**). This is again likely due to the weak Gal4 driver, as the amount of SoxN protein detected in these Tm4s was an order of magnitude lower than in wild-type Tm6 (**Fig. S5A**, insets). Thus, we could engineer predictable switches of type identity between all 4 Tm neurons guided solely by a code of sustained transcription factors.

**Figure 3:**
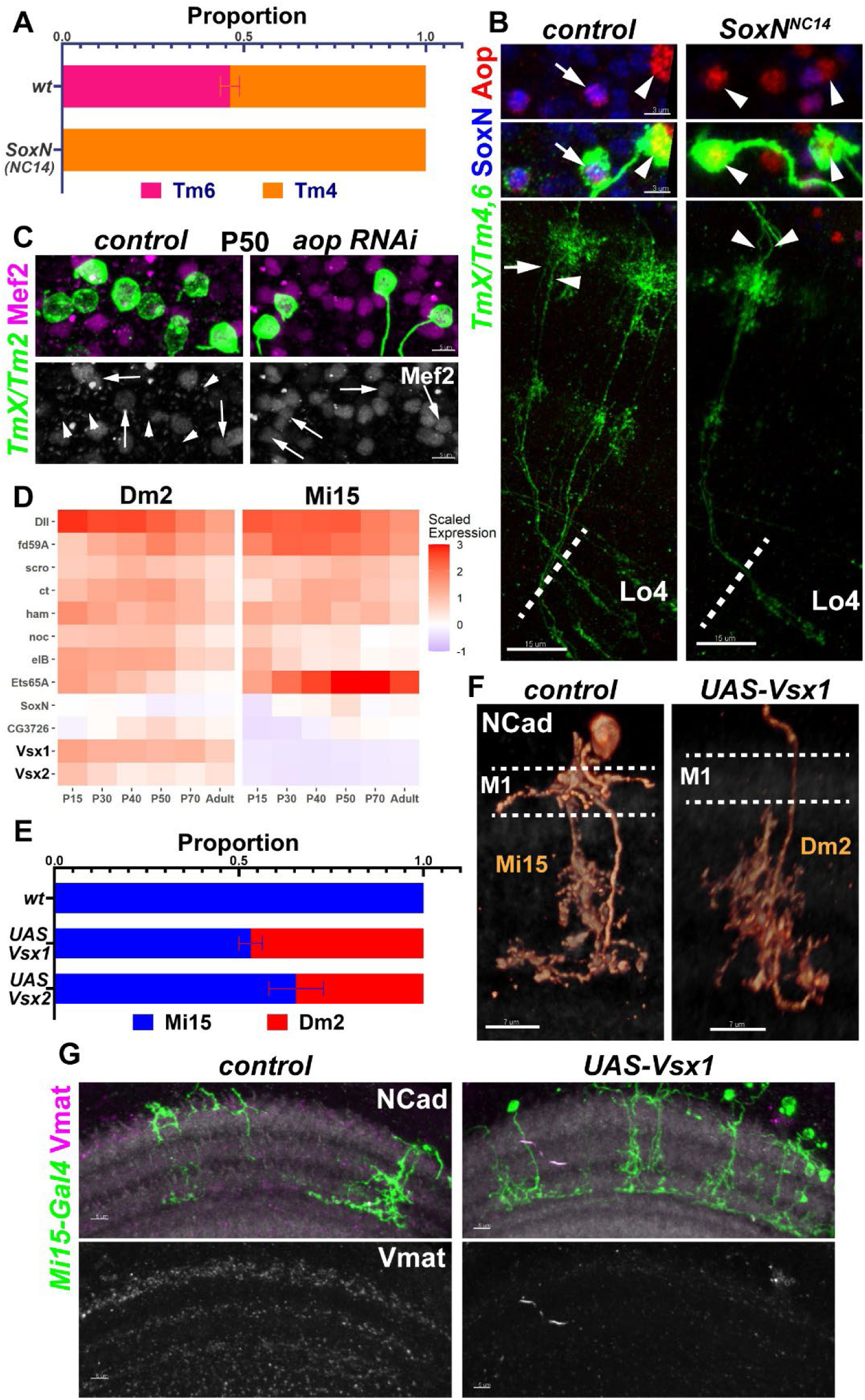
Selector TFs jointly control developmental and functional neuronal type identity. **A-B,** FRT40A (control) and *SoxN^NC^*^14^ MARCM clones labeled with *TmX/Tm4,6-Gal4* and CD4tdGFP in adult brains. **A,** Quantification of B, based on Aop-only (Tm4) and Aop+SoxN (Tm6) neurons. n= 225 neurons in 9 brains (control) and 258 neurons in 10 brains (SoxN^NC^^14^). p<0.0001 for change in Tm6 proportion. In addition, no neurons with Tm6 morphology were observed in the mutant clones. **B,** Max projections with co-stainings of anti-SoxN (blue) and anti-Aop (red), displaying the neurites (bottom) and the cell bodies (top, arrowheads) of the same two neurons occupying the same column. Arrow points to a Tm6 and arrowheads to Tm4s. **C,** *TmX/Tm2-Gal4* driving *aop* RNAi (n=6 brains) and UAS-CD4tdGFP (flip-out). Maximum projections of cell bodies in P50 medulla cortex with co-staining of anti-Mef2 (magenta), which is also shown in single channel (bottom). Arrows point to labeled neurons that express Mef2, and arrowheads indicate those that do not. **D,** Developmental (scaled) expression patterns of all genes that are candidate selectors in either of the displayed cell types (Fig. S1A), based on^5^. Note that scaling was performed over the entire atlas at each stage, thus the values are comparable between the cell types, but not stages. **E,** Quantification of (F) and Fig. S5D. Cells labeled by *Mi15(R76F01)-Gal4* were identified based on their morphology in each condition. n= 104 neurons in 11 brains (wt control), 108 neurons in 11 brains (UAS-Vsx1) and 24 neurons in 4 brains (UAS-Vsx2). p<0.0001 for change in Dm2 proportions in both conditions. Error bars denote SEM. **F-G,** *Mi15-Gal4* driving UAS-Vsx1 and UAS-CD4tdGFP (flip-out). **F,** 3D reconstructions of GFP expression for representative adult neurons in each condition (see Fig. 1A), with co-staining of anti-NCad (white). Dashed lines mark the M1 layer where Mi15 arborizes but Dm2 does not. Also note that Mi15 has two descending branches while Dm2 has one. **G,** Max projections of adult optic lobes with co-stainings of anti-NCad (white) and anti-Vmat (magenta) which is also shown in single channel (bottom). n= 4 (control) and 7 (UAS-Vsx1) brains. Scale bars: 15 μm (B), 5 μm (C,G), 7 μm (F).

Combined, our results suggest that Tm4 is the default fate among these Tm neurons, which is overridden by *Drgx* in Tm1, *pdm3* in Tm2 and *SoxN* in Tm6. *aop* is expressed in both Tm4 and Tm6, but it is repressed by *Drgx* in Tm1 and by *pdm3* in Tm2. To address whether Aop also functions as a selector, we generated *aop* null MARCM clones^47^ and also performed RNAi knock-down using *TmX/Tm4,6-Gal4*; in both cases, the driver was turned off (**Fig. S5B- C**). Instead using *TmX/Tm2-Gal4* to express *aop* RNAi, we observed that all Tm neurons at P50 (when this driver is normally expressed in all 4 Tms) expressed the Tm1/2 marker Mef2 (**Fig. 3C**), indicating that *aop* is necessary for Tm4 and Tm6 identity. However, we could not determine the exact fate of these neurons i.e., eliminated or transformed to Tm1 or Tm2, as both were labeled by the driver.

To further validate the selector TF concept for terminal features of neurons, we sought other neuronal types whose selector codes differ only by a few genes, and also use different neurotransmitters. Dm2 and Mi15 are both cholinergic (like the Tm neurons), but Mi15 are also the only aminergic neurons in the optic lobe^48^, expressing the vesicular monoamine transporter (*Vmat*). They both express the candidate selector TFs *Dll*, *fd59A*, *scro*, *ct*, *ham*, *noc*, *eIB* and *Ets65A*, but *Vsx1* and *Vsx2* are specific to Dm2 and are the only TFs that continuously distinguish the two cell types (**Fig. 3D**). We ectopically expressed *Vsx1* and *Vsx2* using an Mi15-specific (early) driver. Ectopic expression of either *Vsx1* or *Vsx2* in Mi15s was sufficient to convert them to Dm2 morphology (**Fig. 3E-F**, **Fig. S5D**, see **Fig. 1A**). *Vsx1* and *Vsx2* could function redundantly due to their sequence similarity, or they could cross-activate each other’s expression. However, we did not observe Vsx2 protein in Mi15s ectopically expressing Vsx1 (**Fig. S5E**), suggesting redundancy. In addition, using an antibody against Vmat protein^49^, we observed a dramatic reduction in Vmat levels in the medulla upon *Vsx1* overexpression in Mi15 neurons (**Fig. 3G**). The remaining expression is likely from glia or other aminergic neurons in the central brain that send projections to the optic lobe. Taken together, our results show that Vsx genes function as selector TFs in Dm2, controlling both morphological and functional features such as its neurotransmitter.

In summary, we present a comprehensive validation of the selector TF hypothesis outside of *C. elegans*. Our findings strongly support that neuronal identity is primarily encoded by a code of TFs that are continuously maintained in each cell type, from their initial specification to adulthood. These TFs do not seem to specialize as functional units such as morphology or adult terminal features, but rather control all type-specific gene expression throughout differentiation. Critically, we have shown that predictable and complete *in vivo* conversions from one neuronal type to another can be engineered based on this code. To our knowledge, these results represent the first examples of targeted manipulation of neuronal type identity guided solely by high-throughput sequencing data.

### Receptor tyrosine kinase signaling is required to stabilize the Tm selector TF network

Even though scRNA-seq data indicate that the mRNA of the Tm4/6-specific selector TF *aop* could be found in all 4 Tm neurons up to P40 (**Fig. S5F**), Aop protein was not expressed in Tm2 nuclei already at P30 (**Fig. S5G**). This could be explained by a well-known post-translational regulatory mechanism: Aop is exported from the nucleus and degraded after phosphorylation by MAPK^50^. This regulation is essential for the proper specification of R7 photoreceptors in the developing eye through receptor tyrosine kinase (RTK) signaling^51^. We could not detect phosphorylated MAPK in Tm4/6 that strongly express Aop, but P-MAPK was present in Tm1/2 where Aop protein could sometimes only be observed as a ‘ring’ outside the nucleus at P30 (**Fig. S5H**, arrows), suggestive of nuclear export. This suggests that Drgx and Pdm3 initially repress Aop protein indirectly in Tm1 and Tm2 by rendering them sensitive to RTK signaling. This raises the intriguing possibility that neuronal fates could be further sculpted in postmitotic neurons by external signals.

After P40, *aop* mRNA is also downregulated in Tm1 and Tm2; this coincides with *Mef2* upregulation in these cell types (**Fig. S5F**), which we have shown is controlled by Drgx and Pdm3 (**Figs. 1F, 2C**). Knocking-down *Mef2* using *TmX/Tm2-Gal4* resulted in a very rare (1/38 neurons observed) conversion of Tm2 into Tm4 in adults, and 18% of Tm neurons observed were morphologically unrecognizable (**Fig. 4A,D**). Aop could be detected in some of these cells, but only outside the nucleus (**Fig. 4E**), similar to wild-type Tm1/2 at P30 (**Fig. S5H**), before Mef2 activation. This indicates that *aop* was transcriptionally de-repressed without Mef2, but its posttranslational repression through MAPK remained intact, even in adult brains. Therefore, *aop* appears to be downregulated in Tm1/2 through two independent mechanisms (**Fig. 4H**): Degradation of the protein through MAPK at all stages, and transcriptional suppression by *Mef2* after P40. Drgx and Pdm3 could control the first mechanism by regulating the expression of receptors or components of the Ras signaling cascade. Consistently, RTKs genes such as InR, Ror and Alk were differentially expressed between Tm1/2 and Tm4/6 clusters at P30 (**Fig. S5I**).

**Figure 4:**
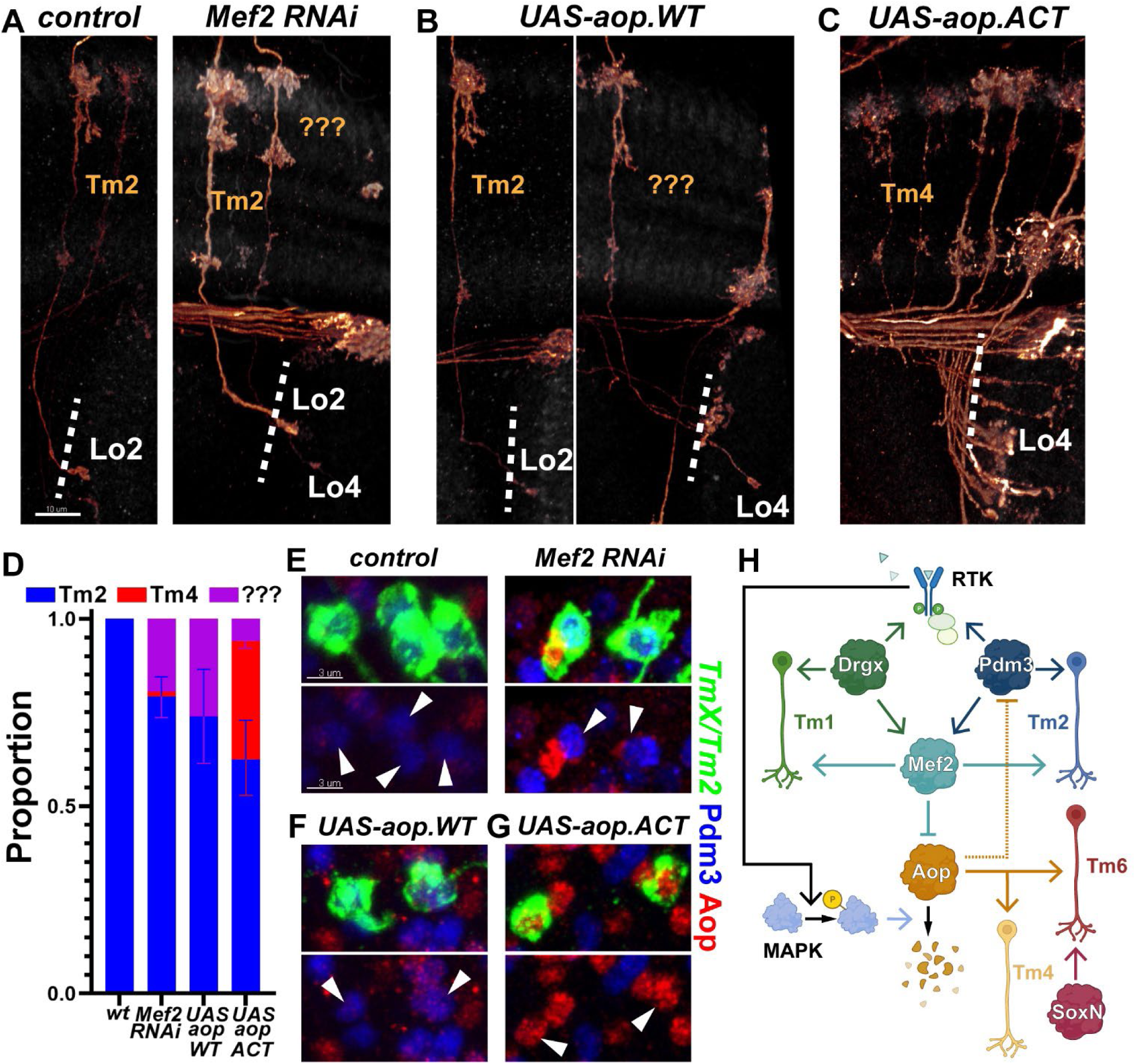
Receptor tyrosine kinase signaling is required to stabilize the Tm selector TF regulatory network. **A-G,** *TmX/Tm2-Gal4* driving UAS-CD4tdGFP (flip-out) and Mef2 RNAi (A,E), UAS-aop.WT (B,F) or UAS-aop.ACT (C,G). **A-C**, 3D reconstructions of GFP expression for the representative adult neurons in each condition (see Fig. 1A), with co-staining of anti-NCad. “???” marks neurons that typically target to Lo4 but could not be recognized as any known optic lobe neuron based on their morphology. Dashed lines mark the border of lobula neuropil. **D,** Quantification of A-C. n= 54 neurons in 6 brains (control), 38 neurons in 8 brains (Mef2 RNAi), 16 neurons in 6 brains (aop.WT) and 75 neurons in 5 brains (aop.ACT). p=0.03 for change in the proportion of unrecognizable neurons with Mef2 RNAi. p=0.02 for change in the proportion of unrecognizable neurons with UAS-aop.WT. p=0.005 for change in the Tm4 proportion with UAS-aop.ACT. Error bars denote SEM. **E-G,** Same as (A-C) with max. projections of cell bodies with co-stainings of anti-Pdm3 (blue) and anti-Aop (red). Arrowheads point to cells labeled with GFP. Scale bars: 10 μm (A-C) and 3 μm (E-G). **H,** Summary of the experimentally validated regulatory interactions between Drgx, Pdm3, Mef2 and Aop in Tm neurons. Negative regulation of Pdm3 by Aop (dashed line) is only applicable when Aop cannot be degraded through the MAPK pathway. RTK: receptor tyrosine kinase.

As *pdm3* and *Drgx* both negatively regulate Aop expression (**Figs. 1G, 2C**), we asked if the opposite was also true. Overexpression of wild-type *aop* with *TmX/Tm2-Gal4* did not convert Tm2 to either Tm4 or Tm6, but 25% of Tm neurons were morphologically unrecognizable (**Fig. 4A,D**), similar to those observed with *Mef2* RNAi where *aop* transcription was derepressed. These neurons still maintained Pdm3 (**Fig. 4F**) whose co-expression with Aop might create a confused state. The signal for the ectopic Aop protein was weak (**Fig. 4F**), suggesting that it was being degraded by the active MAPK pathway in Tm2. We therefore overexpressed a constitutively active form of Aop (*aop.ACT*) that cannot be phosphorylated and degraded^50^. In these brains, 40% of Tm2s were converted to Tm4s (**Fig. 4C-D**) and they had lost Pdm3 expression (**Fig. 4G**). This indicates that while Aop can suppress *pdm3* and promote Tm4 fate (**Fig. 4H**, dashed arrow), this regulation is not relevant with wild-type *aop* due to RTK signaling which ensures that *pdm3* acts upstream.

The apparent destabilization of the fate choice between Tm2 and Tm4 with the postmitotic expression of *aop.ACT* (but not with *aop.*WT) implies that neuronal identity remains dependent on the signaling conditions even after the initial specification events. Similar mechanisms in organisms with larger and more complex brains could be exploited to further diversify the neurons generated from the same stem cell pool with a common identity, but then migrate to distinct brain regions where different signals might be available^52, 53^.

### Decoding neuronal gene regulation through computational network inference

Cell type identity encoded by selector TFs represents only one aspect of neuronal gene regulation. Neuronal transcriptomes are extremely dynamic throughout differentiation, reflecting the sequential developmental steps that neurons traverse: axon guidance and layer targeting, synapse formation, and electrical activity^5, 10^. Importantly, the sustained expression of selector TFs implies that these regulators can be genetically separable from the temporal changes, which are generally controlled by external signals during neuronal maturation^54–56^. For instance, we previously noted that the dramatic diversification of neuronal transcriptomes in the optic lobe just prior to synaptogenesis is preceded by a global upregulation of ecdysone-responsive TFs at P40^5^. This is consistent with a large peak in the hormone titers around P35^57^, as well as the enrichment of EcR motifs in enhancers that become accessible at this stage^27^. This transcriptional timing mechanism has recently been genetically dissected in lamina neurons^14^. It is therefore critical to understand how these two top-level regulatory programs, *i.e.* the identity and developmental state, interact to combinatorially determine the expression of downstream genes in each neuron and at each developmental time to instruct its specific morphology, connectivity and function.

We implemented the Inferelator 3.0 framework^58^ to build gene regulatory network (GRN) models with the goal of gaining a more complete understanding of the regulatory programs employed by developing optic lobe neurons. A key feature of this method is its use of *Transcription Factor Activity* (TFA) that allows Inferelator to estimate the underlying activity of each TF using the expression levels of its known targets from prior information (‘priors’, **Fig. 5A**). Calculating TFA before fitting a linear regression model to infer a GRN circumvents the issue that mRNA levels are not always a good substitute for a TF’s latent activity^59, 60^, which may vary with post-transcriptional modifications (as exemplified by the case of Aop discussed above) or the presence of cofactors.

**Figure 5:**
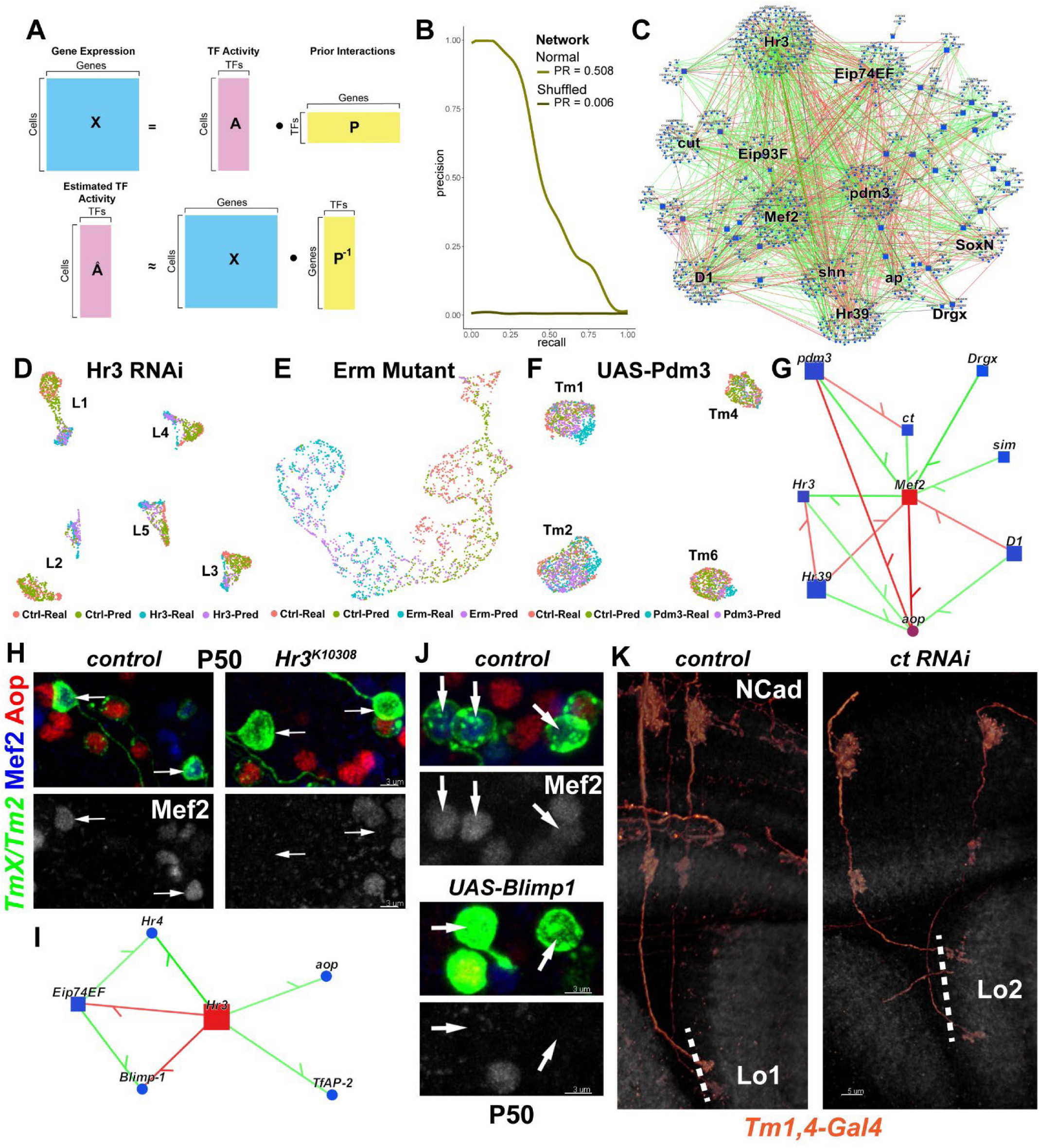
Computational inference of gene regulatory networks and experimental validations. **A,** Transcription factor activity (TFA) calculation. Gene expression in each cell is assumed to be a linear product of latent TF activities in that cell and the connectivity matrix (prior) between TFs and their targets (top). TFA is estimated as the dot product of the expression matrix and the pseudoinverse of the prior matrix (bottom, see Methods). **B**, AUPR curves for the multitask Tm network produced using the MergedDA prior (see also Fig. S6A), and the corresponding negative control network produced using the shuffled prior. **C,** Visualization of the network in (B), displaying all interactions with a minimum of 80% confidence (combined) and variance explained of 1%. Top 10 TFs that had the highest number of target genes in the network above these thresholds were highlighted. SoxN (ranked 13^th^) and Drgx (ranked 24^th^) were not among the top 10 but were nevertheless highlighted as validated selector TFs. **D-F,** scRNA-seq datasets of (**D**) L1-5 neurons at P48 expressing Hr3 RNAi^14^, (**E**) simulated single L3 neurons (Fig. S6D) at P40 mutant for Erm^32^, (**F**) Tm1/2/4/6 neurons at P50 overexpressing UAS- pdm3.short (Fig. 1). Both the original (“Real”) and predicted (“Pred”) (see Fig. S6C) transcriptomes are shown, and the cells are colored according to this status, as well as their condition of origin. UMAPs were calculated using 30 PCs (D,F) or 3 PCs (E) on the integrated (between the real and predicted datasets) gene expression. Note that the cells align correctly according to their cell-type and condition of origin after integration (see Fig. S6E-G for pre- integration UMAPs). **G,** Network visualization displaying all TFs predicted to regulate Mef2 or aop with a min. confidence of 95% in the Tm network, and all inferred interactions between the displayed genes above that threshold. Green: positive, red: negative regulation as determined independently from inference, based on Pearson correlations of mRNA expression between the gene pairs, shading indicates the strength of correlation. **H,** FRT42D (control) and *Hr3^K^*^10308^ MARCM clones labeled with *TmX/Tm2-Gal4* and CD4tdGFP in P50 medulla cortex (maximum projection), with co-stainings of anti-Mef2 (blue) and anti-Aop (red). Arrows point to Aop^-^ Tm neurons that should normally be Mef2^+^ (n=6 brains). **I,** Same as (G), displaying all TFs predicted to be regulated by Hr3 (confidence>95%). **J,** *TmX/Tm2-Gal4* driving UAS-CD4tdGFP (flip-out) and UAS-Blimp1 (n=5 brains). Max. projection same as (H). **K,** *Tm1,4-Gal4* driving UAS- CD4tdGFP (flip-out) and *ct* RNAi (n= 40 Tm1 neurons in 6 brains). 3D reconstructions of GFP expression for the representative adult neurons in each condition with co-staining of anti-NCad (white). Dashed lines mark the border of lobula neuropil. Scale bars: 3 μm (H-J) and 5 μm (K).

We modeled GRNs for two distinct groups of cell types: the four types of Tm neurons investigated in this paper, as well as the five types of lamina monopolar neurons (L1-5), which are among the most intensely studied cell types in the optic lobe and provide a useful benchmark for our models. We focused on the early-mid pupal (P24-50) stages when optic lobe neurons acquire most of their specific morphological features and begin to form synapses. For inference, we used single-cell transcriptomes from both our atlas^5^, at stages P30, P40 and P50, as well as one generated by another group^61^, at stages P24, P36 and P48. Using the scATAC-seq dataset described above^27^, we constructed priors for each network: First, we identified differentially accessible regions between Tm1/2/4/6 clusters (Tm network) and L1-5 clusters (Lamina network) at P48 (**Fig. S4C**). These regions were scanned and scored within 10kb of every transcription unit for TF binding motifs. These scores were then clustered to retain only the highest confidence targets of each TF for TFA calculation. We tested three different prior generation methods (**Fig. S6** displays benchmarking results for all of them, see Methods for extended discussion), but the “MergedDA” priors were used for all final networks. The networks were then modeled only on the genes that displayed differential expression between either the cell-types or the time-points analyzed (1394 genes for Tm, 1684 for Lamina networks). The network performances were evaluated against the respective priors using the following metrics: area under the precision/recall curves (AUPR), Matthews correlation coefficient (MCC), F1 scores, and the distribution of variance explained by the inferred edges (**Fig. S6A-B**, see Methods for detailed description). We additionally compared the performance of each network to negative control networks that were built with shuffled priors (**Fig. 5B**, **Fig. S6A-B)**. **Figure 5C** displays the entire GRN inferred from the Tm neurons, highlighting the top 10 TFs with the highest number of targets predicted, which includes *Mef2*, the selector TFs *pdm3*, *cut* and *ap*, as well as the ecdysone-responsive TFs *Hr3*, *Hr39*, *Eip74EF* and *Eip93F*.

The scarcity of ‘gold standard’ networks presents a challenge when benchmarking inferred GRNs in complex multicellular organisms. To assess the predictive power of our models, we exploited available RNA-seq datasets collected from perturbed neurons. For lamina neurons, we used two relevant datasets: a scRNA-seq study of all 5 lamina neurons with knock- down of *Hr3* at P48^14^; and a bulk RNA-seq study at P40 of FACSed L3 neurons mutant for Erm^32^, a selector TF of these neurons (**Fig. S1A**). First, we calculated the TFA in each cell in the Hr3 knock-down dataset with the same Lamina prior used for inference. We then applied matrix multiplication (dot product) between the estimated TFA and the learned weights between TFs and targets (betas) to generate a predicted expression matrix (**Fig. S6C**). It is important to emphasize that *Hr3* knock-down creates novel regulatory states in all 5 lamina neurons that were never seen by the inference algorithm, as the betas were determined from completely independent, wild-type datasets discussed above. Despite these challenges, after Seurat integration^62^, the real and predicted transcriptomes aligned nearly perfectly according to their cell-type (L1-5) and condition (control vs. knock-down) of origin (**Fig. 5D**). Since the Erm dataset was generated with bulk sequencing, we first simulated single-cell transcriptomes from each of the 5 replicates of control vs. mutant libraries (**Fig. S6D**). The predicted transcriptomes (using the same Lamina prior and betas) were again well-aligned with the corresponding condition in the real dataset (**Fig. 5E**). However, the UMAP visualizations of the unintegrated transcriptomes (**Fig. S6E-G**) showed that the differences between control and perturbed conditions were much smaller in predicted clusters compared to the real ones. We analyzed this by determining the differentially expressed genes (DEGs) between the control and perturbed conditions for both real and predicted clusters. For each cell type, we calculated the ratio of the number of DEGs in predicted clusters that were also DEGs between the corresponding real clusters (precision), and the ratio of correctly predicted DEGs to all DEGs in the real clusters (recall). We found that the predicted transcriptomes recapitulated only a small proportion (5-20%) of the real DEGs (low recall), but the predicted DEGs were mostly (>50%) correct (high precision). To similarly benchmark the Tm networks, we used the scRNA-seq dataset we acquired from the Tm neurons overexpressing *pdm3* at P50 (**Fig. 1H**). Like the lamina datasets, the real and predicted transcriptomes showed good concordance (**Fig. 5F**), with even higher precision (70-90%) as well as relatively better recall (30-40%) of DEGs between most clusters (**Fig. S6J**).

In summary, by combining single-cell gene expression and chromatin accessibility datasets, we were able to build predictive computational models of GRNs in optic lobe neurons. Our benchmarks suggest that the interactions learned by our models are largely accurate, though they represent only a snippet of the true underlying GRNs. We thereby propose that these models will be useful to understand how selector TFs may control downstream effectors that implement various aspects of neuronal differentiation during development.

### Selector TFs interact with ecdysone signaling to regulate downstream targets

We interrogated our GRN models to understand how selector TFs may interact with other regulatory programs such as those controlled by ecdysone. Specifically, we aimed to understand how these different regulatory programs can together control the spatiotemporally specific expression of other TFs. We focused on the specific subnetwork upstream of *Mef2* and *aop* (**Fig. 5G**) and observed that nearly all the regulatory relationships we experimentally validated in the previous sections (**Fig. 4H**) were also captured by our GRN model: positive regulation of *Mef2* by *Drgx* and *pdm3*, and negative regulation of *aop* by *pdm3* and *Mef2*. As discussed above, Drgx and Pdm3, while required, are not sufficient to activate *Mef2* expression, which does not occur until P40 despite the continuous expression of the selector TFs. Ecdysone-responsive *Hr3* emerged as an attractive candidate for this temporal trigger: despite being predicted as upstream of *aop* but downstream of *Mef2* in the model, it is more likely to be regulating both, as its activation around P30 precedes *Mef2* (**Fig. S5F**, note that the Inferelator does not consider the temporal order of the stages used). We tested this prediction by generating *Hr3* mutant MARCM clones using *TmX/Tm2-Gal4*. In adult brains, Mef2 expression was not affected and Tm2 neurons appeared morphologically normal (**Fig. S7A**). However, at P50, Mef2 could not be detected in mutant clones (**Fig. 5H**), indicating that *Mef2* expression was delayed, but not abolished in *Hr3* mutants. This indicates that there are redundant temporal mechanisms regulating *Mef2* expression and/or that *Hr3* acts indirectly to control *Mef2*. Among the TFs that were predicted to be downstream of *Hr3* in our GRN model (**Fig. 5I**), *Hr4*, *Eip74EF* and *Blimp-1* were all shown to be regulated by *Hr3* in lamina neurons^14^. *Blimp-1* is normally expressed at high levels during early pupal stages. It starts being downregulated at P40 and disappears completely by P50 (**Fig. S5F**). Interestingly, upon *Hr3* knock-down, this downregulation is delayed but not abolished in all 5 lamina neurons^14^. We thus asked whether *Blimp-1* functions as a repressor of *Mef2*, through which *Hr3* might act. Indeed, overexpression of *Blimp-1* using *TmX/Tm2-Gal4* also repressed Mef2 at P50 (**Fig. 5J**), validating this prediction. This phenotype could not be assessed in adults, as the flies that express *UAS-Blimp-1* did not eclose, even when the expression was restricted to pupal stages using *Gal80^ts^*. Therefore, by experimentally validating the predictions of our computational models, we described how the combinatorial action of the selector TFs *Drgx*/*Pdm3* and the ecdysone-responsive TFs *Hr3*/*Blimp-1* enables *Mef2* to be expressed specifically in Tm1 and Tm2 neurons, and only after P40.

Another inferred edge in this sub-network (**Fig. 5G**), the negative regulation of *cut* (*ct*) by *pdm3*, was consistent with the lower levels of *ct* expression in Tm2s compared to other Tm neurons (**Fig. S7B**). Since different Cut expression levels in larval da sensory neurons regulate the size of their dendritic arborizations^63^, we asked whether this difference in *ct* levels among Tm neurons is functionally significant. Indeed, we found that 73% of Tm1s targeted to Lo2 layer instead of Lo1 with *ct* RNAi knock-down (**Fig. 5K**). Even though *ct* was a candidate selector TF due to its continuous expression in all 4 Tm neurons (**Fig. 1B**), these results show that different levels of its expression (as modified by other selectors) control a specific feature during brain wiring; thus it also functions as an effector controlling a sub-routine. However, overexpression of *ct* in Tm2 neurons did not lead to their axons projecting to Lo1 instead of Lo2 (**Fig. S7C**). Therefore, downregulation of *ct* levels by *pdm3* in Tm2s appears to be sufficient but not necessary for targeting to Lo2, suggesting that there are redundant mechanisms allowing them to arborize in this layer.

Lastly, we inspected the inferred GRN model of the Tm neurons (**Fig. 5C**) to assess if different types of TFs specialize on different types of targets. We previously reported that TFs and cell-surface proteins (CSPs) are overrepresented in DEGs between optic lobe neurons; CSPs are particularly upregulated at P40-50^5^. This is consistent with the proposed importance of these proteins in synaptic specificity^64^, and the fact that synaptogenesis in the optic lobe commences around these stages^65^. Thus expectedly, CSPs that may be involved in cell-cell recognition^66^ were strongly enriched among the target genes included in the filtered Tm network (**Fig. S7D**, “All”). We then examined the percentage of these genes among the inferred targets of the top 10 TFs in the network with the highest number of targets. We did not see a clear bias for any TF class to regulate more or fewer CSPs than others (**Fig. S7D**). However, the two selector TFs included in the top 10 list, *ap* and *pdm3*, had among the highest ratios of inferred targets that were also TFs (ranked 1^st^ and 4^th^, **Fig. S7D**). While anecdotal, this finding suggests that the broad changes caused by selector TF perturbations could be partially due to a higher likelihood of selectors to regulate other TFs. We also performed GO enrichment analysis on the predicted targets of the top 3 TFs in the network: ecdysone TF Hr3, selector TF Pdm3 and effector TF Mef2. The same general terms were enriched for all of them: ion channels, cell adhesion and signaling molecules (**Fig. S7E**). This similarity was not due to these TFs regulating highly overlapping sets of genes. While the three TFs had some common targets as expected, most of the predicted targets (54-62%) for each of these TFs were not among the targets of the other two. These results further highlight the combinatorial nature of neuronal gene regulation and they are consistent with other recent findings that most targets of the ecdysone-responsive TFs are cell-type specific^14^, despite the uniform expression of these TFs in all neurons.

Altogether, we presented here a genetic cascade from a temporal TF that initially specifies neuronal fates in progenitors to the selector TFs that initiate and maintain these fates in postmitotic neurons. We then relied on computational GRN models to describe how these selectors control the effector genes downstream and used these models to genetically dissect the temporal regulation of *Mef2* expression by ecdysone signaling.

## Discussion

### Selector TFs: Strengths and limitations

We set out to test a simple but powerful idea: Neuronal type identity is primarily encoded by unique and sustained combinations of transcription factors in each cell type. Originally put forth in *C. elegans*, the terminal selector hypothesis has been supported by overwhelming evidence in worms, and a few TFs that appear to fit the definition have been found in other systems as well (reviewed in^67^). But it remained unclear how generally applicable this logic could be for encoding neuronal type identity in more complex brains. We refer to these genes as “selector TFs”, as it is clear that they do not only encode a “terminal” state, but also orchestrate a series of dynamic developmental programs. The original emphasis on the terminal (which may be interpreted as adult) features of neurons, led others to conclude that several TFs that appear to be continuously expressed and can induce morphological switches of neuronal identity should not be considered terminal selectors because they do not affect the neurotransmitter identity of those motoneurons that are all glutamatergic^25^. *Drgx*, *pdm3*, *SoxN* and *aop* studied here also do not affect neurotransmitter identity, as all 4 Tm neurons are cholinergic^5^ which must be controlled by other selector genes, such as *ap*^68^. It is nevertheless clear that selector TFs together regulate all type-specific gene expression among those neurons, as shown by scRNA-seq. Our results do not rule out the presence of TFs that only function to control morphology or connectivity, but such genes would likely be transiently expressed during development and be downstream of selector TFs in the regulatory hierarchy. Lastly, unlike the “classical” selector genes that were proposed to be clonally maintained to specify entire tissues or organs^69^, neuronal selectors function post-mitotically to commit individual cell types to specific fates.

A chief strength of the selector TF concept is that tTFs from neuroblasts and/or Notch effectors from GMCs can directly relay information about specification through the activation of selectors in newly born neurons. Our results suggest that this is the case from the tTF *Klu* to the selector TF *Drgx*. Previous studies in the optic lobe have found additional genetic links between known tTFs and neuronal TFs which we now propose are candidate selector TFs based on their sustained expression in the respective neurons (**Fig. S1A**). The tTF Hth regulates Bsh (selector of Mi1), and Ey regulates Drf, a selector TF for ∼9 medulla neurons^11, 30, 43^. The tTF Slp1 regulates the widely expressed selectors Toy and Sox102F, and the tTF D regulates Ets65A^11, 70^, which is a common selector of Mi15 and Dm2 (among others). tTF Hbn also regulates Toy, and other selectors including Tj and Otd; and the tTF Opa regulates TfAP-2^10^, a selector TF shared by all four Tm neurons we studied here. Finally, all Notch^ON^ neurons express the selector TF Ap^11^, which is required for the cholinergic identity of most of those neurons^68^. In addition, the tTFs Hth, Ey, D, Scro and Erm are maintained (or re-activated) in some of the neurons produced from those temporal windows (but not all, except for Hth)^10^ and continue to function as selectors (**Fig. S1A**). It is not surprising that the genetic program that specifies neuronal identities in progenitors is intricately linked to the one that initiates and maintains them in neurons. But we still know little about how the combined action of temporal, spatial and Notch patterning activates a unique set of selectors in every neuronal type, and subsequently how the selector combination enacts precise gene batteries over the course of development. Integrative analysis of single-cell transcriptomic and epigenomic datasets will allow us to better understand these mechanisms in the future.

The apparent conservation of this regulatory logic in both *C. elegans* and *Drosophila*, whose last common ancestor lived over 600 million years ago^71^, makes it likely that the selector TF concept will also be useful to understand and manipulate the neuronal diversity of the mammalian brain. This could have large implications for the emerging field of cell replacement therapy. The current protocols to generate specific neuronal types *in vitro* mostly rely on recapitulating the signaling conditions that the relevant progenitors normally go through and/or providing lineage-specific TFs to pluripotent stem cells^72, 73^. These require extensive prior knowledge about their development that is only available for a few cell types. Furthermore, the efficiency of these protocols tends to be low, and the resulting neurons are often heterogenous populations of related cell types. We propose that induction of more ‘generic’ neurons from *in vitro* stem cells using pan-neuronal factors^74^, followed by transfection of specific selector TF cocktails could be an effective alternative to these approaches. The increasing prevalence of developmental scRNA-seq atlases can enable the discovery of these TF combinations based on their sustained expression pattern, as we have achieved here for optic lobe neurons, at high precision and scale. However, our results suggest that the signaling environment in which these transplanted neurons will be placed can also impact their final identity.

There are a few notable limitations of the selector TF framework that should be taken into consideration. First, we and others have previously reported that a few neuronal types have distinct transcriptomes during development but then converge to a common state in adult brains^5, 75^. These are generally very similar (sub)types that only differ in their connectivity, thus the TFs that encode their differences do not need to be maintained after their wiring is complete. Second, the earliest events of neuronal differentiation, such as neuropil targeting which in the optic lobe happens immediately after their terminal division can potentially be under the control of factors that are only transiently expressed during the first few hours of a neuron’s life. Such genes may be directly regulated by tTFs and/or Notch and not the selector TFs. As these decisions are likely to be irreversible once completed due to physical constraints of anatomy, this could obstruct complete morphological conversions between neurons that target to entirely different neuropils. Lastly, selector TFs can be post-transcriptionally regulated by, for instance, RNA binding proteins Imp and Syp that are widely utilized in *Drosophila* brain to generate neuronal diversity^76, 77^. This could complicate the identification of correct selector TF combinations from RNA-seq data alone, as we have done in this study.

### Understanding brain wiring through gene regulatory networks

The constancy of selector TF expression implies that their regulation is independent from the temporal transitions that occur through pupal development as the neurons differentiate and form circuits. This provides a unique opportunity to understand developmental gene regulatory networks (GRNs) where the selector TFs can be confidently placed at the top of the regulatory hierarchy, enabling easier dissection of the downstream mechanisms. Accurate computational inference of GRNs describing relationships between TFs and their target genes has been a long-standing goal of systems biology. GRN reconstruction has benefited immensely from the recent advancements in single-cell genomics with a myriad of different methods becoming available (reviewed in^78, 79^). As models derived from mRNA expression alone can only provide correlative predictions, state-of-the-art methods typically combine gene expression data with cis-regulatory information such as chromatin accessibility to infer causal interactions^27, 52^. However, most inference algorithms still rely on correlating mRNA levels between target genes and their potential regulators, which may not reflect the true latent activity of TFs in every cell type. The Inferelator approach we use in this work avoids this issue by first estimating this underlying activity using a short list of high-quality targets for each TF. These prior connectivity matrices can be derived from binding assays (e.g. ChIP-seq) or chromatin accessibility (e.g. ATAC-seq), as we have done here. Importantly, unlike the other methods that combine single-cell expression and accessibility datasets for GRN inference^27, 80, 81^, we used these priors derived from scATAC-seq for TFA calculation only, and thus could also discover interactions that are not necessarily present in the prior, some of which may be indirect. Our benchmarks suggest that the learned models can accurately generalize to novel regulatory states created by the perturbation of specific TFs. Additionally, our experimental validations that showed how the ecdysone signaling co-operates with selector TFs to regulate the spatiotemporally restricted expression of the effector TF *Mef2* highlights the power of these models in discovering novel regulatory interactions. L3 selector TF Erm was also recently shown to interact with EcR to regulate a variety of targets in these neurons^14^.

Both this work and other recent efforts to decipher gene regulation in the fly brain^27^ have now made it possible to study the molecular mechanisms of synaptic specificity within the framework of gene regulatory mechanisms that encode neuronal type identity. We propose a “top-down” approach where selector TFs that cause broad changes in neuronal fates are identified first, followed by the dissection of downstream mechanisms aided by GRN modeling. Perhaps the most promising targets are effector TFs, that still regulate many other genes but have more limited functions (sub-routines) than the selector TFs. We found that low levels of *cut* in Tm2, downstream of the selector TF *pdm3*, specifically controls the axon targeting choice to Lo1 vs. Lo2 layers. Nevertheless, the current models still have significant limitations, imposed mainly by the quality of priors that can be derived from chromatin accessibility. In addition, the current scATAC-seq atlas of the fly brain^27^ has considerably lower cell-type resolution than the comparable scRNA-seq atlases, especially during early development. Single-cell multiome studies that simultaneously profile gene expression and chromatin accessibility in the same cells could remove many of these limitations in the near future.

## Supporting information

FACS Gating Strategy

## Acknowledgements

We would like to thank all members of the Desplan and Bonneau Labs, Oliver Hobert, Richard Mann and Denis Jabaudon for helpful discussions. We thank Chris Doe, Stein Aerts, Justin Blau, Esteban Mazzoni, Robin Hiesinger, Nikos Konstantinides, David Chen, Ryan Loker and Sromana Mukherjee for critical reading of the manuscript. We further thank David Krantz, Ilaria Rebay, Cheng-Ting Chien, Wesley Grueber and Jens Rister for reagents. We would like to extend special thanks to Jasper Janssens and Stein Aerts for sharing the scATAC-seq data ahead of publication, and to Giuseppe-Antonio Saldi for help with network inference. This work was supported by NIH grants EY13010 and EY017916 to C.D. and by R01HD096770, R01CA229235 and RM1HG011014 to R.B and by Simons Foundation. M.N.O. was a Leon Levy Neuroscience Fellow and is supported by NINDS K99NS125117. C.S.G. is supported by the NSF Award 1922658 to NYU CDS. I.H. was supported by an HFSP postdoctoral fellowship (LT000757/2017-L) and by the Kimmel Center for Stem Cell Biology Senior Postdoctoral Fellowship.

## Author Contributions

M.N.O., R.B. and C.D. conceived the project. M.N.O. designed all experiments. C.S.G. developed and optimized the Inferelator workflows. M.N.O., I.H. and M.S. performed the experiments. M.N.O. and C.S.G. analyzed the data. M.N.O., C.S.G. and C.D. wrote the manuscript. All authors edited the manuscript.

## Data Availability

Raw and processed scRNA-seq data are publicly accessible on GEO: GSE199734.

---

**Figure S1:**
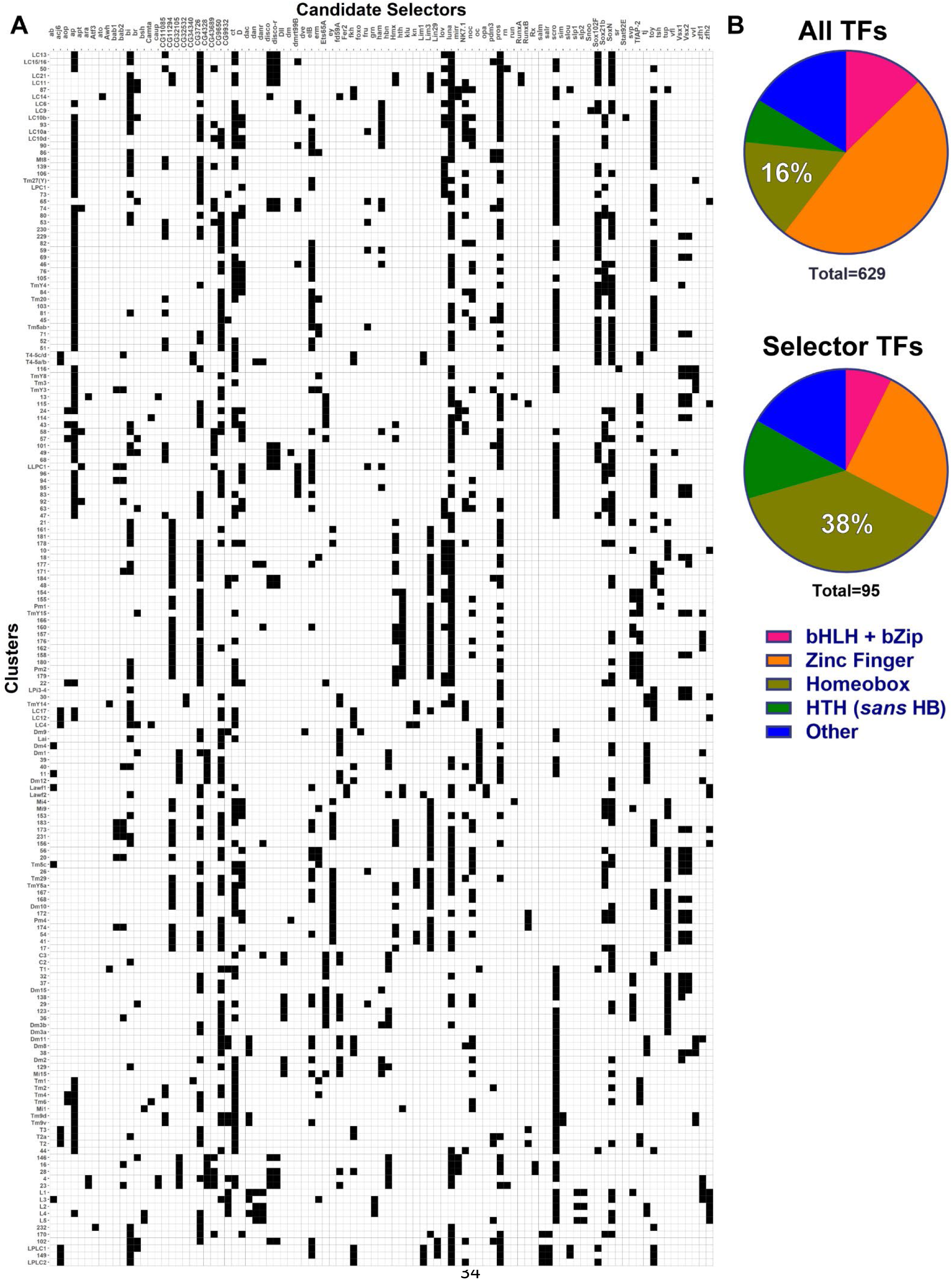
Candidate selector TFs of the optic lobe neurons. **A,** Differentially expressed TFs (columns) found to be continuously expressed in each optic lobe neuron (rows) throughout its development according to^5^, see Methods and Table S1. **B,** Pie charts displaying the proportions of major TF classes within all TFs in the *Drosophila* genome (top), and the TFs found to be a candidate selector for at least one cell type in A. Basic domain, zinc finger, homeobox, helix-turn-helix (excluding homeobox) categories are represented, with the other TF classes merged in “Other”.

**Figure S2:**
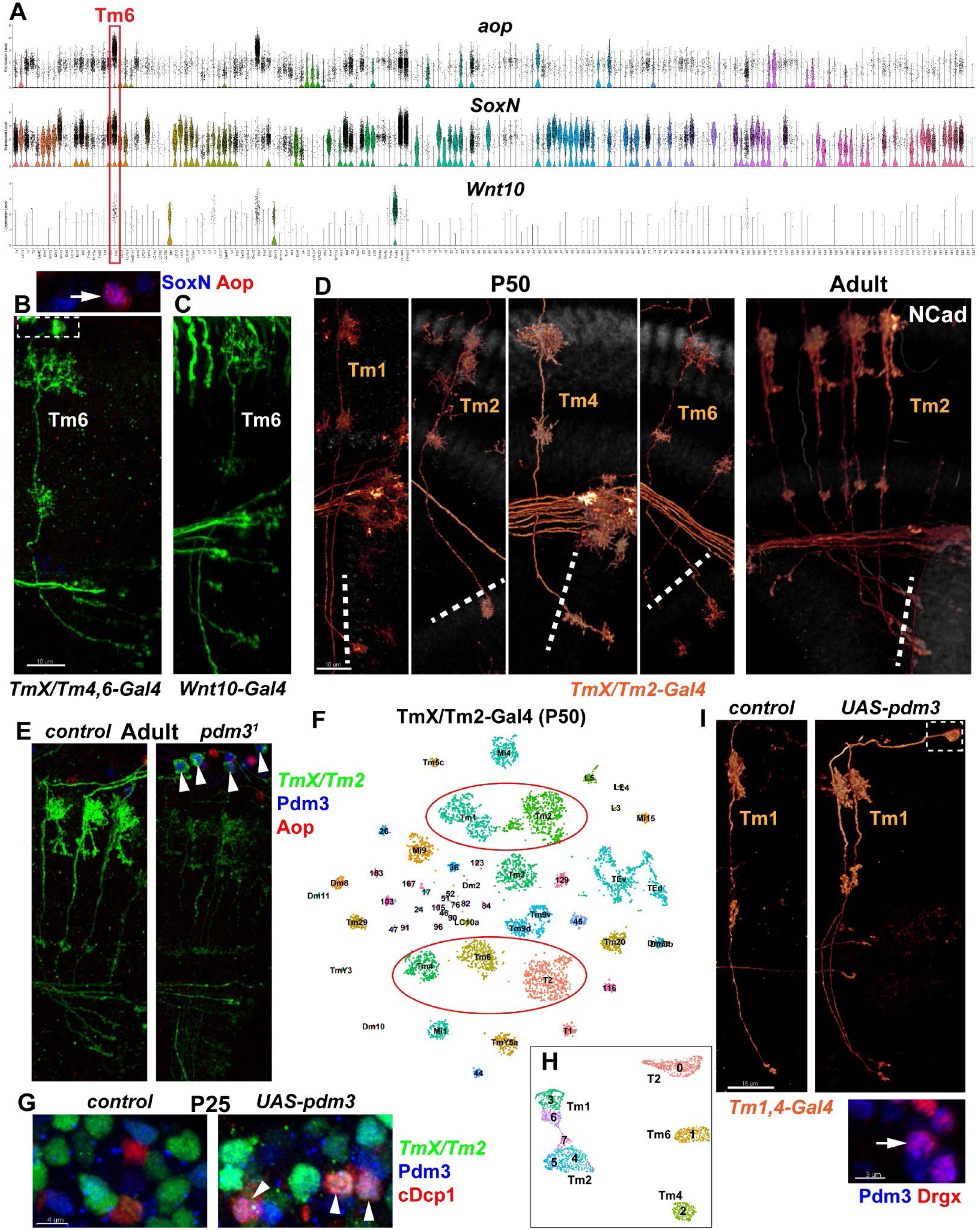
Tm6 annotation and *pdm3* as the selector TF of Tm2 neurons (Supplement to Fig. 1) **A,** Violin plots displaying the log-normalized expression of *aop*, *SoxN* and *Wnt10* in all adult optic lobe neurons^5^. Cluster #62 (red rectangle) exclusively expresses all 3 genes. **B,** Max. projections of *TmX/Tm4,6-Gal4* (left) or *Wnt10-Gal4* (right) driving UAS-CD4tdGFP (flip-out), displaying an adult Tm6 neuron (see Fig. 1A) with co-stainings of anti-SoxN (blue) and anti- Aop (red) for the inset in B. Arrow points to the nucleus of the neuron displayed below. **D,** 3D reconstructions of *TmX/Tm2-Gal4* driving UAS-CD4tdGFP (flip-out) at P50 and adult optic lobes with co-staining of anti-NCad (white), displaying representative Tm neurons. Dashed lines mark the border of lobula neuropil. **E,** FRT40A (control) and *pdm3*^1^ MARCM clones labeled with *TmX/Tm2-Gal4* and CD4tdGFP in adult optic lobes (maximum projection), with co-stainings of anti-Pdm3 (blue) and anti-Aop (red). Only Tm neurons observed in the mutants still expressed Pdm3 (arrowheads) and were labeled very weakly with GFP, suggesting this is leaky expression from MARCM (incomplete Gal80 suppression) rather than mutant clones (n=10 brains). **F,** scRNA-seq of FACSed neurons from P50 optic lobes, *TmX/Tm2-Gal4* driving nuclear GFP (Stinger) and UAS-pdm3.short. tSNE was calculated using 50 principal components. Cells are colored and labeled according to neural network^5^ classifications. Red ellipses mark the cells subset for further analysis (Fig. 1H-O). **G,** Max. projections of P25 medulla cortex, with *TmX/Tm2-Gal4* driving UAS-Stinger (green) and UAS-pdm3.short (right), with co-stainings of anti-Pdm3 (blue) and anti-cleaved Dcp1 (red) marking apoptotic cells. Arrowheads indicate the nuclei with triple labeling (n= 5 brains per condition, quantified in the main text). **H,** Same as in Fig. 1J, with cells colored and labeled according to unsupervised clustering (as in 1K). **I,** 3D reconstructions of *Tm1,4-Gal4* driving UAS-CD4tdGFP (flip-out) and UAS-pdm3.short (n=6 brains), displaying representative adult Tm1 neurons. Inset shows a max. projection of the indicated cell body (arrow) with co-stainings of anti-Pdm3 (blue) and anti- Drgx (red). Scale bars: 10 μm (B-E), 4 μm (G), 15 μm (I) and 3 μm (I, inset).

**Figure S3:**
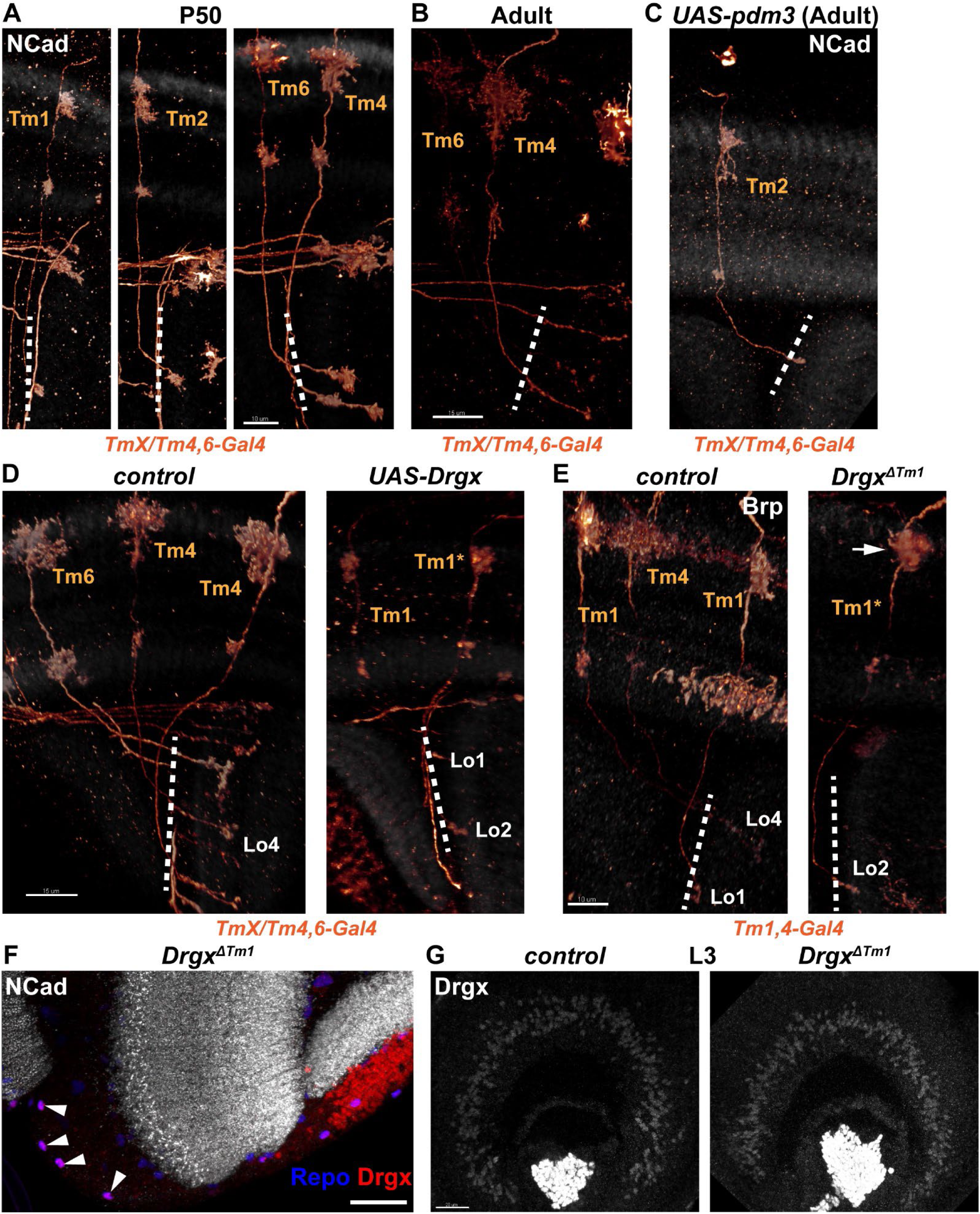
Drgx as the selector TF of Tm1 neurons (Supplement to Fig. 2) **A-C,** 3D reconstructions of *TmX/Tm4,6-Gal4* driving UAS-CD4tdGFP (flip-out) and UAS- pdm3.short (**C** only) with co-staining of anti-NCad (white), displaying representative Tm neurons in P50 (**A**) and adult (**B-C**) optic lobes. Note that generally (n=4 brains) no Tm neurons were labeled by the driver in (C), except in the one brain shown here where a single Tm2 neuron was recovered. **D,** 3D reconstructions of *TmX/Tm4,6-Gal4* driving UAS-CD4tdGFP (flip-out) and UAS-Drgx with co-staining of anti-NCad (white), displaying representative adult Tm neurons (same as in Fig. 2A). The Tm1 labeled with an asterisk abnormally targets to Lo2 layer instead of Lo1. Quantified in Fig. 2B. **E,** 3D reconstructions of adult neurons with *Tm1,4-Gal4* driving UAS-CD4tdGFP (flip-out) in the background of heterozygous (control) or homozygous *Drgx^ΔTm1^* allele (same as in Fig. 2G), with co-staining of anti-Brp (white). Quantified in Fig. 2H. The Tm1 labeled with an asterisk abnormally targets to Lo2 and has a disrupted dendritic arbor (arrow). **F,** Same as in Fig. 2G with co-staining of anti-NCad (white), anti-Repo (blue) and anti- Drgx (red), showing that the only remaining Drgx^+^ cells in the medulla cortex (arrowheads) of *Drgx^ΔTm1^* mutants are glia (n=3 brains). **G,** Max. projections of L3 optic lobes showing anti-Drgx staining in w^11^^18^ (control, n=5 brains) and homozygous *Drgx^ΔTm1^* mutants (n=7 brains). Scale bars: 10 μm (A, E-F), 15 μm (B-D) and 20 μm (G). Dashed lines mark the border of lobula neuropil.

**Figure S4:**
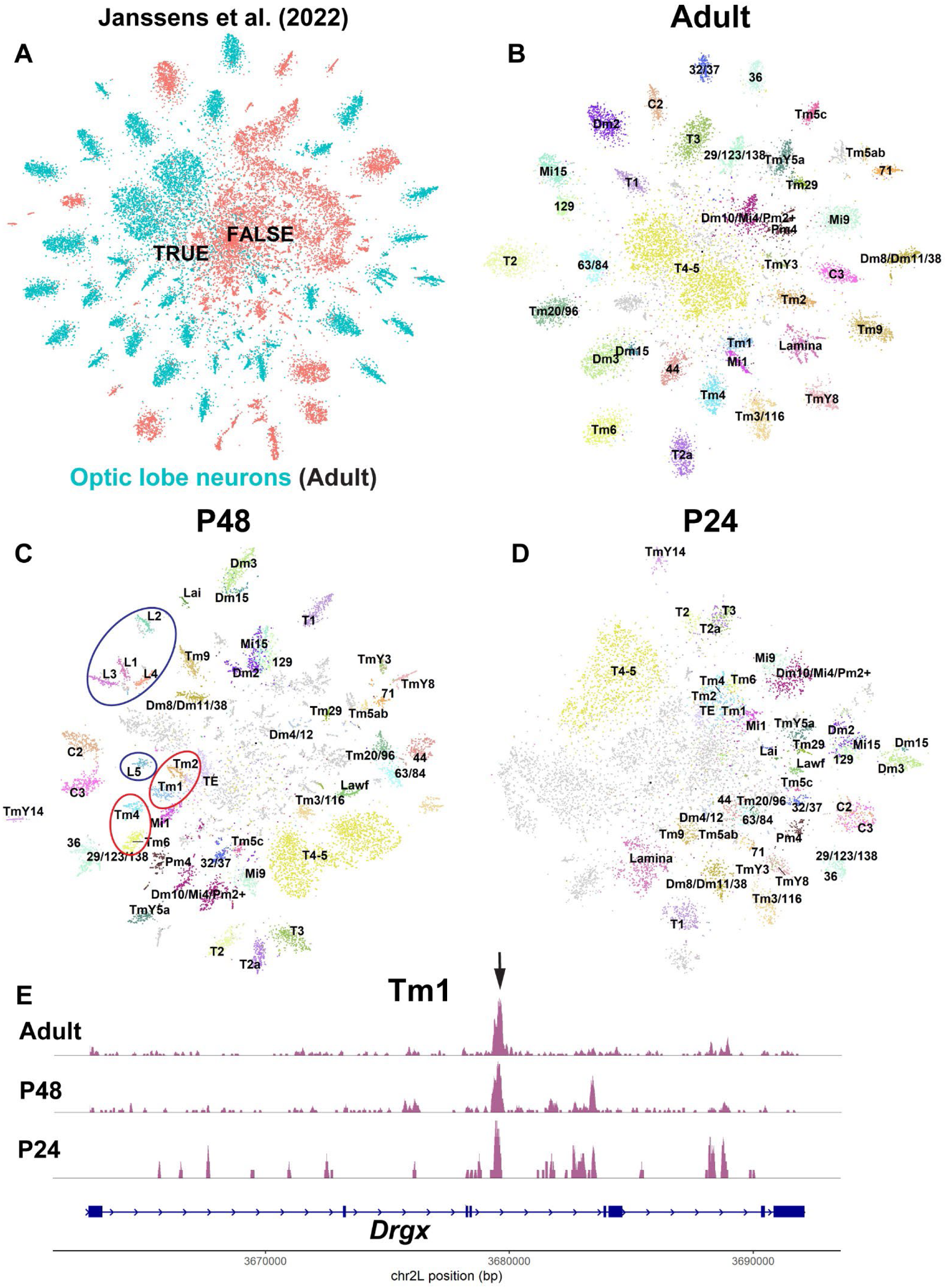
Annotation of optic lobe neurons in scATAC-seq datasets. **A-C,** Clusters corresponding to the optic lobe neurons in Adult (A), P48 and P24 scATAC-seq experiments^27^ were isolated, clustered and annotated (**B-D**) separately at the indicated stages (see Methods for details). tSNE visualizations were calculated using the top 120 principal components, excluding the first PC. Clusters labeled by numbers are unannotated neurons in the reference scRNA-seq atlas^5^. Clusters that could not be linked to defined (groups of) clusters in the reference atlas were grayed out. Ellipses in (C) indicate the clusters used for building the Inferelator priors for lamina (blue) and Tm (red) neurons for GRN inference. **E,** Aggregated accessibility tracks of *Drgx* locus for the Tm1 cluster, using the TF-IDF normalized scATAC-seq data at the indicated stages (B-D). Arrow points to the Tm1-specific enhancer deleted in (Fig. 2G-I). See also Fig. 2D.

**Figure S5:**
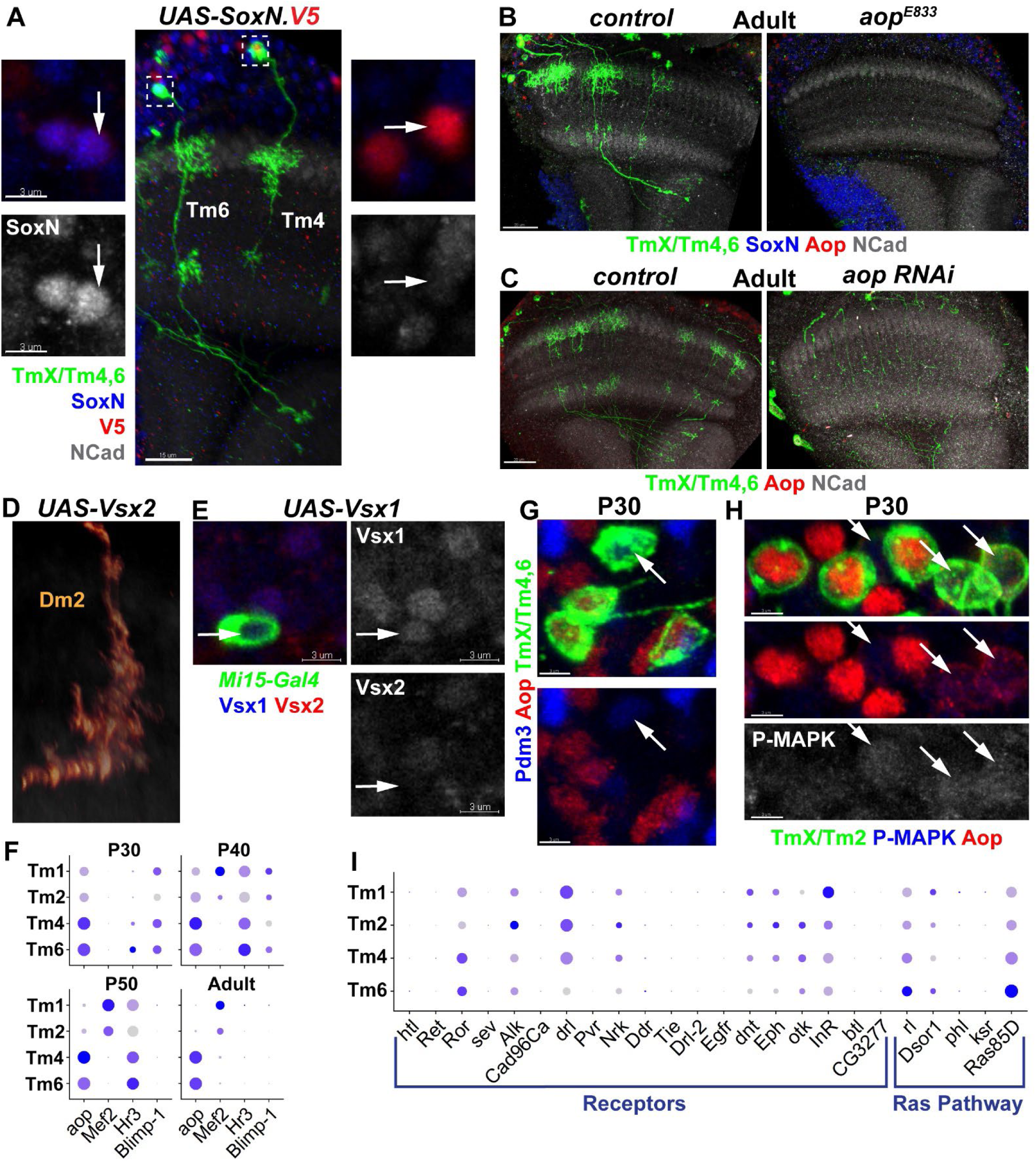
Additional selector TFs of optic lobe neurons and regulation of the Tm selector network by RTK signaling (Supplement to Figs. 3 and 4) **A,** *TmX/Tm4,6-Gal4* driving UAS-SoxN.V5 and UAS-CD4tdGFP (flip-out, n=3 brains). Max. projection of an adult optic lobe with co-stainings of anti-NCad (white), anti-SoxN (blue) and anti-V5 (red). Insets show the indicated cell bodies (arrows) with anti-SoxN also shown in single channel (bottom). **B,** Max. projections of adult optic lobes in FRT40A (control) and *aop^E8^*^33^ MARCM clones (n=3 brains) labeled with *TmX/Tm4,6-Gal4* and CD4tdGFP, with co-stainings of anti-NCad (white), anti-SoxN (blue) and anti-Aop (red). **C,** Max. projections of adult optic lobes with *TmX/Tm4,6-Gal4* driving UAS-CD4tdGFP (flip-out) and *aop* RNAi (n=8 brains), with co-stainings of anti-NCad (white) and anti-Aop (red). **D,** Same as in Fig. 3F, with *Mi15-Gal4* driving UAS-Vsx2 instead of Vsx1. Quantified in Fig. 3E. **E,** Same as in Fig. 3G, max. projection of cell bodies with co-stainings of anti-Vsx1 (blue) and anti-Vsx2 (red) that are also shown in single channel (n=3 brains). Note that Vsx1/2 are almost always co-expressed in the optic lobes except for the Mi15 neurons in this experiment that express Vsx1 ectopically (arrow). **F,I,** Dot plots displaying the log-normalized expression in Tm1/2/4/6 clusters of indicated TFs in the P30, P40, P50 and adult scRNA-seq datasets^5^ (**F**), or of all RTK receptors and Ras signaling pathway components expressed in the optic lobe at P30 (**I**). Diameters of the dots indicate the proportion of cells in each cluster expressing the genes, while the shading is proportional to the level of expression. G, *TmX/Tm4,6-Gal4* driving UAS-CD4tdGFP (flip-out) in P30 medulla cortex (max projection), with co-stainings of anti-Pdm3 (blue) and anti-Aop (red), arrow indicates a GFP^+^Pdm3^+^ Tm2 in which no Aop protein can be detected (n=3 brains)**. H**, *TmX/Tm2-Gal4* driving UAS-CD4tdGFP (flip-out) in P30 medulla cortex (max projection), with co-stainings of anti-P-MAPK (blue) and anti-Aop (red). Arrows indicate GFP^+^Aop^-^ Tm1/2 neurons in which PMAPK (single-channel, bottom) can be detected. Note that Aop^+^ neurons visible do not display PMAPK signal (n=5 brains). Scale bars: 15 μm (A), 3 μm (A-insets, E, G, H) and 20 μm (B-C).

**Figure S6:**
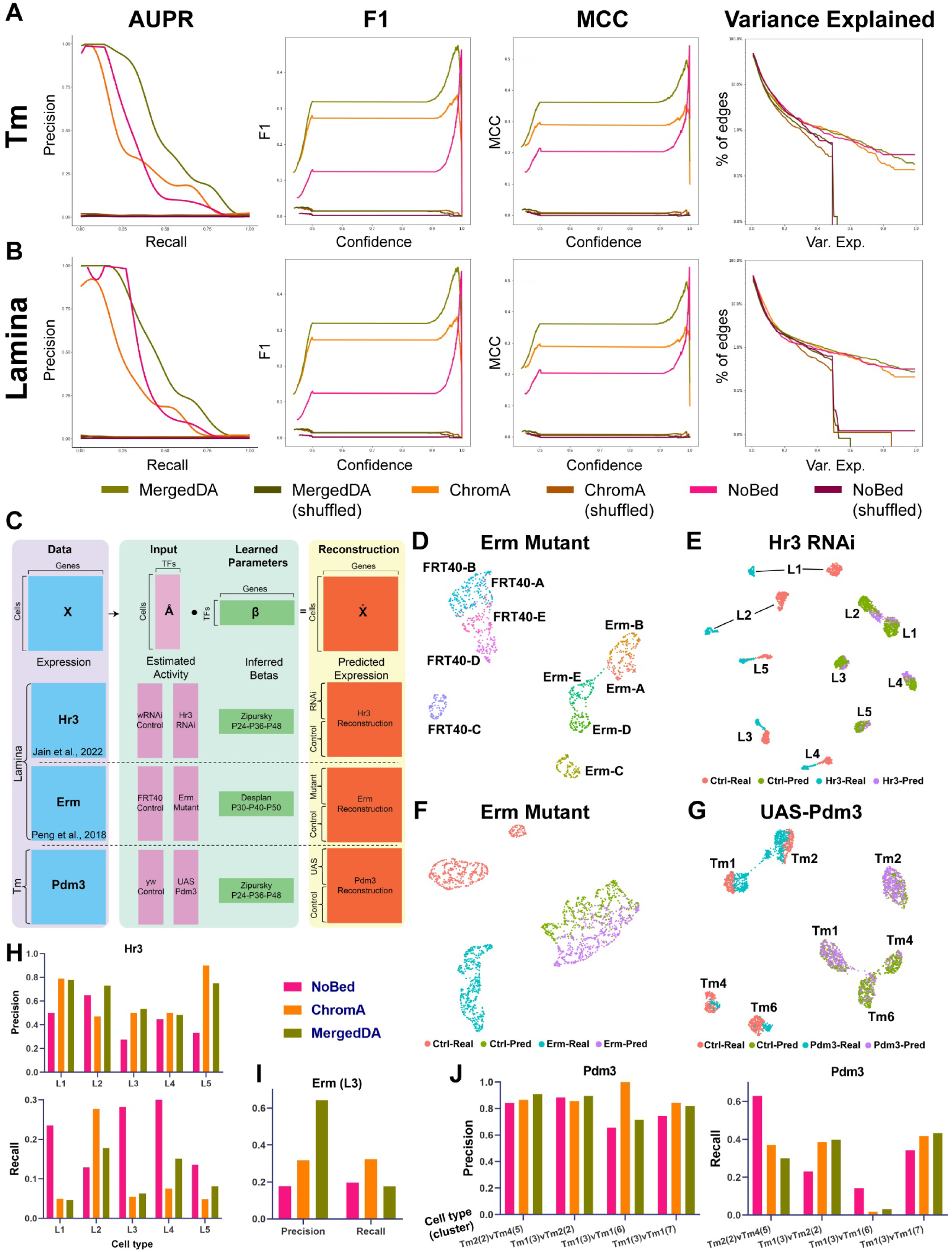
Computational inference and benchmarking of GRNs in optic lobe neurons (Supplement to Fig. 5) **A-B,** Curves of Area under Precision and Recall (AUPR), F1 scores, Mathews Correlation Coefficient (MCC) and variance explained by the edges inferred (multitask) from (**A**) Tm1/2/4/6 and (**B**) L1-5 cell types using three different priors for each network, and their shuffled negative controls. See Methods for detailed description of the priors and displayed metrics. **C,** Reconstruction method for predicting gene expression profiles of the RNA-seq experiments shown. TFA is calculated for each cell as described in Fig. 5A using the same priors for the respective lamina (for Hr3 and Erm) and Tm (for Pdm3) network inference. Dot products of these TFA matrices with the learned betas from the indicated inference tasks, i.e., the weights between TFs and their targets as determined by the Inferelator using only the wild-type data, generate a predicted expression matrix matching the original data. **D,** Simulated single-cell transcriptomes (100 per replicate, see Methods) from the Erm bulk RNA-seq experiment at P40^32^. UMAP visualization was calculated using 3 principal components (PCs). **E-G,** scRNA-seq datasets of (**E**) L1-5 neurons at P48 expressing Hr3 RNAi^14^, (**F**) simulated-single Erm mutant L3 neurons from (D), (**G**) Tm1/2/4/6 neurons at P50 overexpressing UAS-pdm3.short (Fig. 1). Both the original (“Real”) and predicted (“Pred”) (based on networks built with the MergedDA prior) transcriptomes are shown, and the cells are colored according to this status, as well as their condition of origin. UMAPs were calculated using 30 PCs (E,G) or 3 PCs (F) on the non-integrated gene expression. See Fig. 5D-F for UMAPs of the same data after Seurat integration was performed between the real and predicted clusters. **H-J,** Comparative analysis of real and predicted differentially expressed genes (DEGs) between the experimental conditions for cell types or clusters shown in E-G (see Fig. 1K for the cluster numbers in J), calculated separately using the networks generated with three different priors. Precision is defined as the ratio of correctly determined DEGs between the conditions in predicted clusters (i.e., those that were also DEGs between the real clusters), and the recall is defined as the ratio of these correctly predicted DEGs to all DEGs between the real clusters.

**Figure S7:**
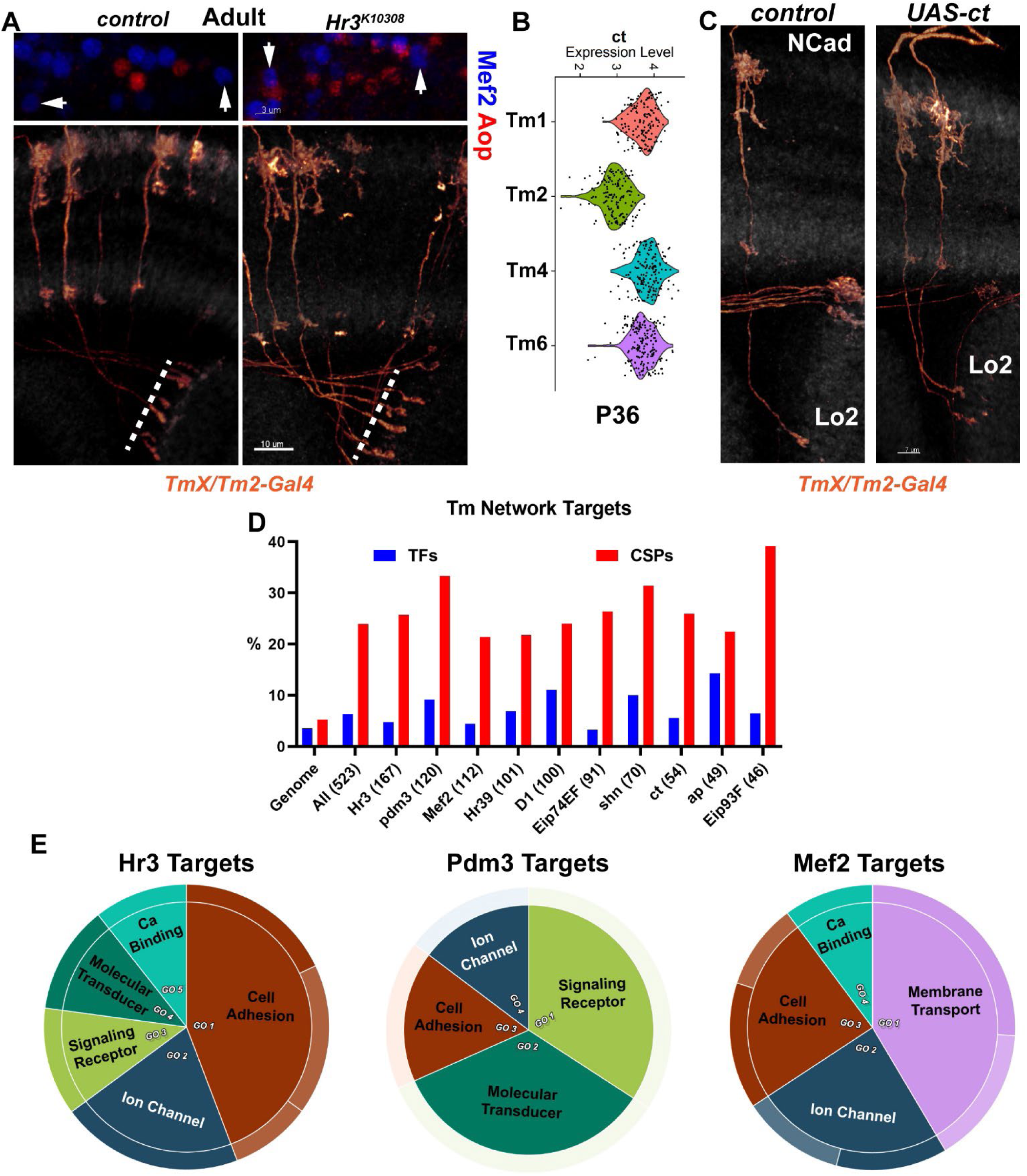
Additional validation experiments for the GRN models of Tm neurons (Supplement to Fig. 5) **A,** FRT42D (control) and *Hr3^K103^*^08^ MARCM clones labeled with *TmX/Tm2-Gal4* and CD4tdGFP showing (bottom) 3D reconstructions of representative adult neurons with co-staining of anti-NCad (white), and the max. projections of cell bodies with co-stainings of anti-Mef2 (blue) and anti-Aop (red). Arrows point to the nuclei of Tm2 neurons shown in bottom panels (n=5 brains). Dashed lines mark the border of lobula neuropil. **B,** Violin plot displaying the log-normalized expression of *ct* in Tm1/2/4/6 clusters in the P36 scRNA-seq dataset^61^. **C,** *TmX/Tm2-Gal4* driving UAS-CD4tdGFP (flip-out) and UAS-ct (n=6 brains). 3D reconstructions of GFP expression for the representative neurons in each condition with co-staining of anti-NCad (white). Scale bars: 10 μm (A-bottom), 3 μm (A-top) and 7 μm (C). **D,** Percentages of genes encoding TFs and cell-surface proteins (CSPs) in the entire genome (17,551 genes), among all target genes included in the filtered GRN model of Tm neurons (the same as in Fig. 5C), and among the predicted targets (in the same model) of each TF shown. The total number of genes considered for each group are indicated in parentheses. **E,** GO enrichment analysis of the predicted targets of Hr3, Pdm3 and Mef2 in the filtered Tm GRN model (same as in D). Molecular Function terms with greater than twofold enrichment were summarized by REVIGO to eliminate redundant terms and group related ones together. Areas within the graphs were determined by the p-values of the terms.

**Table S1**: **Candidate selector TFs of the optic lobe neurons: Source data for Figure S1A.**

**Table S2**: **Differential gene expression analysis of the Tm subclusters in scRNA-seq**

**Table S3: Genotypes of animals used for all experiments in figure panels and the temperature at which each experiment was performed**

**Table S4: Inferelator models learned from the Tm and Lamina neurons**: Produced using the respective MergedDA priors, multitask from the P30-40-50 and P24-36-48 datasets.

**Supplementary Information:** FACS gating strategy.

## Methods

### Key Resources Table

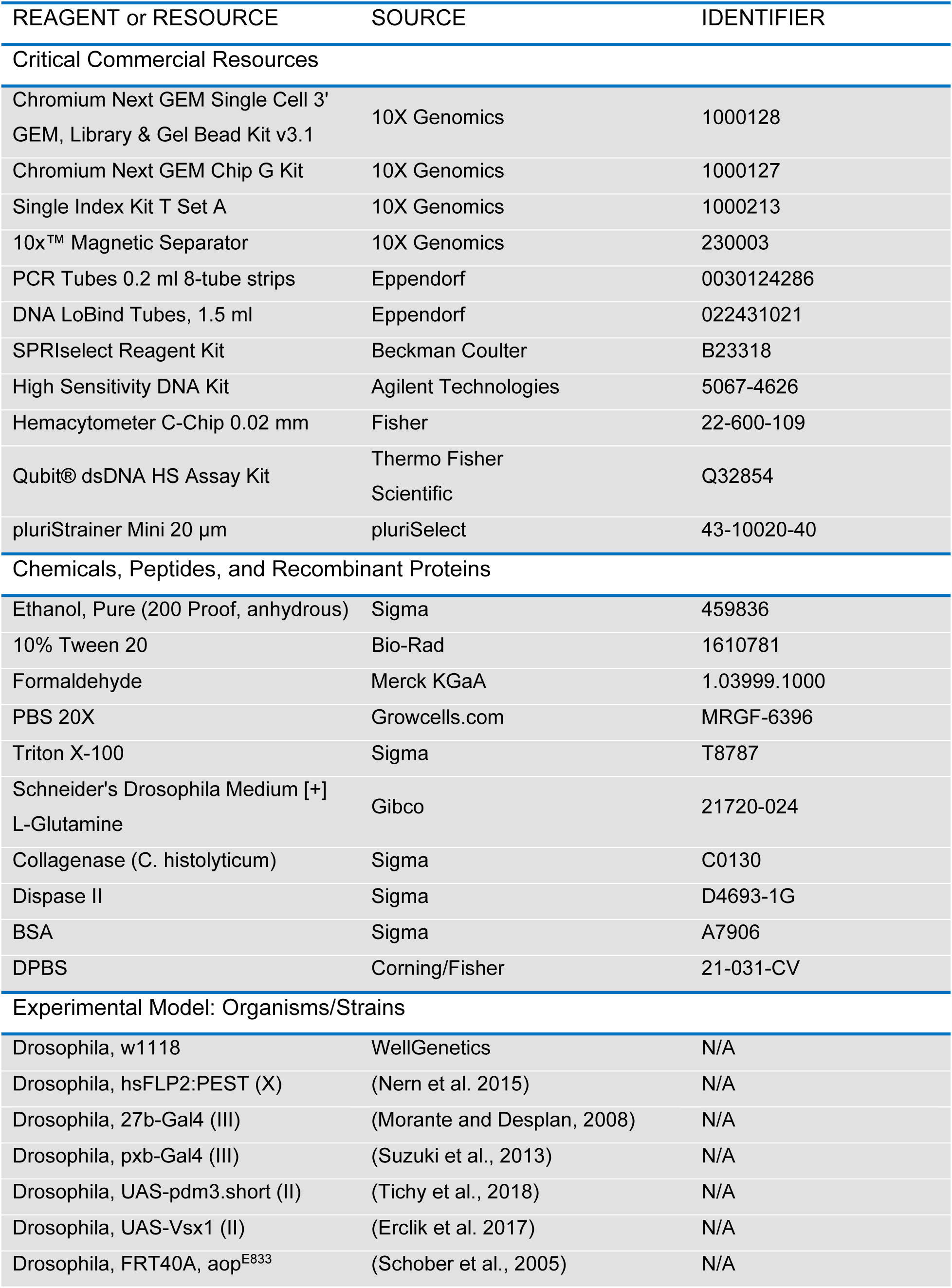
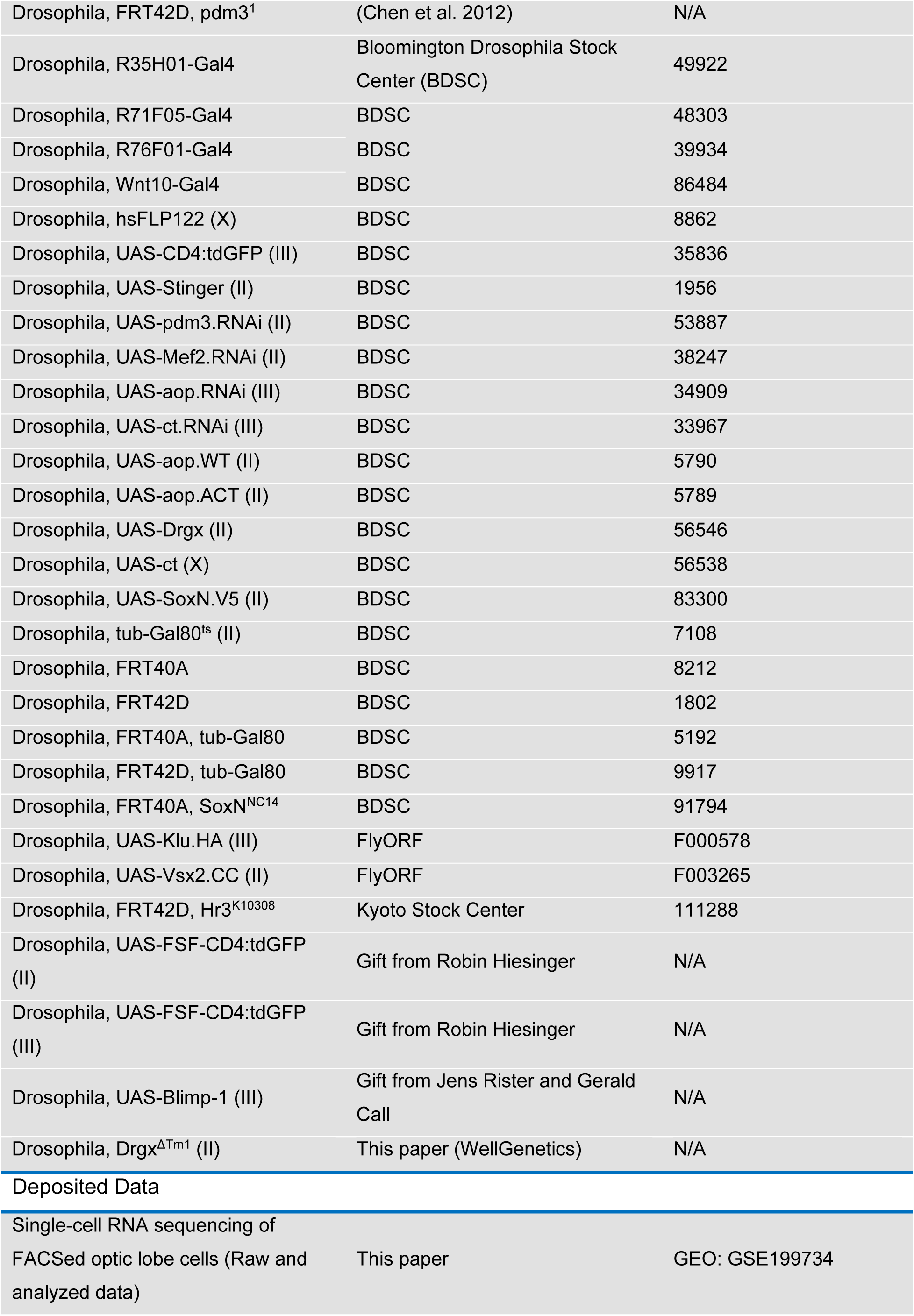
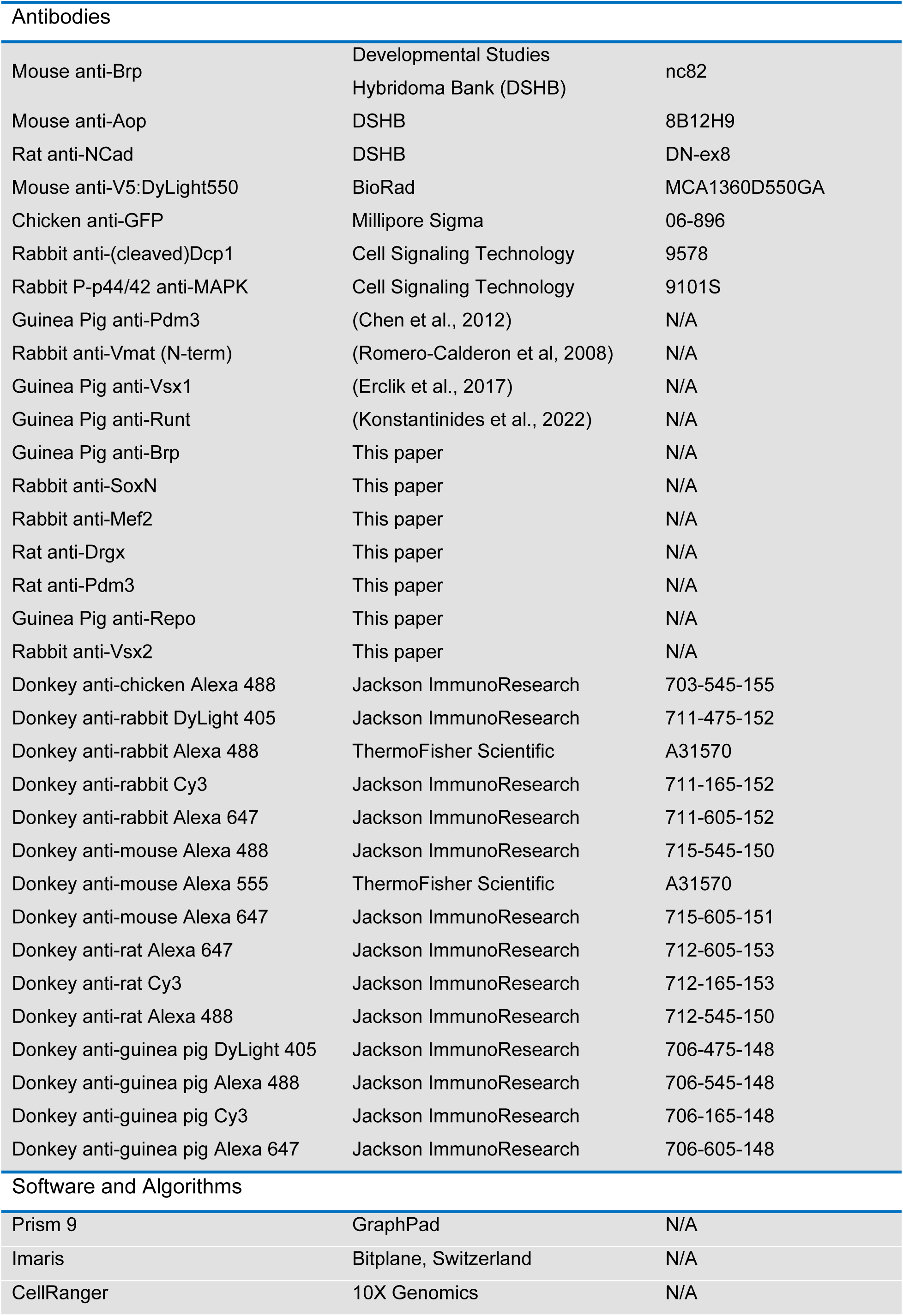
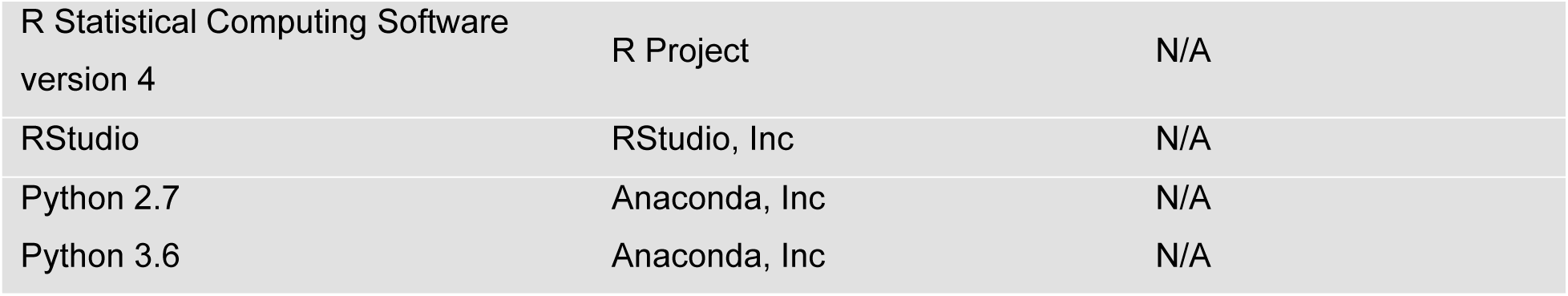

| REAGENT or RESOURCE | SOURCE | IDENTIFIER |
| --- | --- | --- |
| Critical Commercial Resources |  |  |
| Chromium Next GEM Single Cell 3' GEM, Library & Gel Bead Kit v3.1 | 10X Genomics | 1000128 |
| Chromium Next GEM Chip G Kit | 10X Genomics | 1000127 |
| Single Index Kit T Set A | 10X Genomics | 1000213 |
| 10x™ Magnetic Separator | 10X Genomics | 230003 |
| PCR Tubes 0.2 ml 8-tube strips | Eppendorf | 0030124286 |
| DNA LoBind Tubes, 1.5 ml | Eppendorf | 022431021 |
| SPRIselect Reagent Kit | Beckman Coulter | B23318 |
| High Sensitivity DNA Kit | Agilent Technologies | 5067-4626 |
| Hemocytometer C-Chip 0.02 mm | Fisher | 22-600-109 |
| Qubit® dsDNA HS Assay Kit | Thermo Fisher Scientific | Q32854 |
| pluriStrainer Mini 20 µm | pluriSelect | 43-10020-40 |
| Chemicals, Peptides, and Recombinant Proteins |  |  |
| Ethanol, Pure (200 Proof, anhydrous) | Sigma | 459836 |
| 10% Tween 20 | Bio-Rad | 1610781 |
| Formaldehyde | Merck KGaA | 1.03999.1000 |
| PBS 20X | Growcells.com | MRGF-6396 |
| Triton X-100 | Sigma | T8787 |
| Schneider's Drosophila Medium [+]<br>L-Glutamine | Gibco | 21720-024 |
| Collagenase (C. histolyticum) | Sigma | C0130 |
| Dispase II | Sigma | D4693-1G |
| BSA | Sigma | A7906 |
| DPBS | Corning/Fisher | 21-031-CV |
| Experimental Model: Organisms/Strains |  |  |
| Drosophila, w1118 | WellGenetics | N/A |
| Drosophila, hsFLP2:PEST (X) | (Nern et al. 2015) | N/A |
| Drosophila, 27b-Gal4 (III) | (Morante and Desplan, 2008) | N/A |
| Drosophila, pxb-Gal4 (III) | (Suzuki et al., 2013) | N/A |
| Drosophila, UAS-pdm3.short (II) | (Tichy et al., 2018) | N/A |
| Drosophila, UAS-Vsx1 (II) | (Erclik et al. 2017) | N/A |
| Drosophila, FRT40A, aop<sup>E833</sup> | (Schober et al., 2005) | N/A |
| Drosophila, FRT42D, pdm3<sup>1</sup> | (Chen et al. 2012) | N/A |
| Drosophila, R35H01-Gal4 | Bloomington Drosophila Stock Center (BDSC) | 49922 |
| Drosophila, R71F05-Gal4 | BDSC | 48303 |
| Drosophila, R76F01-Gal4 | BDSC | 39934 |
| Drosophila, Wnt10-Gal4 | BDSC | 86484 |
| Drosophila, hsFLP122 (X) | BDSC | 8862 |
| Drosophila, UAS-CD4:tdGFP (III) | BDSC | 35836 |
| Drosophila, UAS-Stinger (II) | BDSC | 1956 |
| Drosophila, UAS-pdm3.RNAi (II) | BDSC | 53887 |
| Drosophila, UAS-Mef2.RNAi (II) | BDSC | 38247 |
| Drosophila, UAS-aop.RNAi (III) | BDSC | 34909 |
| Drosophila, UAS-ct.RNAi (III) | BDSC | 33967 |
| Drosophila, UAS-aop.WT (II) | BDSC | 5790 |
| Drosophila, UAS-aop.ACT (II) | BDSC | 5789 |
| Drosophila, UAS-Drgx (II) | BDSC | 56546 |
| Drosophila, UAS-ct (X) | BDSC | 56538 |
| Drosophila, UAS-SoxN.V5 (II) | BDSC | 83300 |
| Drosophila, tub-Gal80<sup>ts</sup> (II) | BDSC | 7108 |
| Drosophila, FRT40A | BDSC | 8212 |
| Drosophila, FRT42D | BDSC | 1802 |
| Drosophila, FRT40A, tub-Gal80 | BDSC | 5192 |
| Drosophila, FRT42D, tub-Gal80 | BDSC | 9917 |
| Drosophila, FRT40A, SoxN<sup>NC14</sup> | BDSC | 91794 |
| Drosophila, UAS-Klu.HA (III) | FlyORF | F000578 |
| Drosophila, UAS-Vsx2.CC (II) | FlyORF | F003265 |
| Drosophila, FRT42D, Hr3<sup>K10308</sup> | Kyoto Stock Center | 111288 |
| Drosophila, UAS-FSF-CD4:tdGFP (II) | Gift from Robin Hiesinger | N/A |
| Drosophila, UAS-FSF-CD4:tdGFP (III) | Gift from Robin Hiesinger | N/A |
| Drosophila, UAS-Blimp-1 (III) | Gift from Jens Rister and Gerald Call | N/A |
| Drosophila, Drgx<sup>ΔTm1</sup> (II) | This paper (WellGenetics) | N/A |
| Deposited Data |  |  |
| Single-cell RNA sequencing of FACSed optic lobe cells (Raw and analyzed data) | This paper | GEO: GSE199734 |

| Antibodies |  |  |
| --- | --- | --- |
| Mouse anti-Brp | Developmental Studies Hybridoma Bank (DSHB) | nc82 |
| Mouse anti-Aop | DSHB | 8B12H9 |
| Rat anti-NCad | DSHB | DN-ex8 |
| Mouse anti-V5:DyLight550 | BioRad | MCA1360D550GA |
| Chicken anti-GFP | Millipore Sigma | 06-896 |
| Rabbit anti-(cleaved)Dcp1 | Cell Signaling Technology | 9578 |
| Rabbit P-p44/42 anti-MAPK | Cell Signaling Technology | 9101S |
| Guinea Pig anti-Pdm3 | (Chen et al., 2012) | N/A |
| Rabbit anti-Vmat (N-term) | (Romero-Calderon et al, 2008) | N/A |
| Guinea Pig anti-Vsx1 | (Erclik et al., 2017) | N/A |
| Guinea Pig anti-Runt | (Konstantinides et al., 2022) | N/A |
| Guinea Pig anti-Brp | This paper | N/A |
| Rabbit anti-SoxN | This paper | N/A |
| Rabbit anti-Mef2 | This paper | N/A |
| Rat anti-Drgx | This paper | N/A |
| Rat anti-Pdm3 | This paper | N/A |
| Guinea Pig anti-Repo | This paper | N/A |
| Rabbit anti-Vsx2 | This paper | N/A |
| Donkey anti-chicken Alexa 488 | Jackson ImmunoResearch | 703-545-155 |
| Donkey anti-rabbit DyLight 405 | Jackson ImmunoResearch | 711-475-152 |
| Donkey anti-rabbit Alexa 488 | ThermoFisher Scientific | A31570 |
| Donkey anti-rabbit Cy3 | Jackson ImmunoResearch | 711-165-152 |
| Donkey anti-rabbit Alexa 647 | Jackson ImmunoResearch | 711-605-152 |
| Donkey anti-mouse Alexa 488 | Jackson ImmunoResearch | 715-545-150 |
| Donkey anti-mouse Alexa 555 | ThermoFisher Scientific | A31570 |
| Donkey anti-mouse Alexa 647 | Jackson ImmunoResearch | 715-605-151 |
| Donkey anti-rat Alexa 647 | Jackson ImmunoResearch | 712-605-153 |
| Donkey anti-rat Cy3 | Jackson ImmunoResearch | 712-165-153 |
| Donkey anti-rat Alexa 488 | Jackson ImmunoResearch | 712-545-150 |
| Donkey anti-guinea pig DyLight 405 | Jackson ImmunoResearch | 706-475-148 |
| Donkey anti-guinea pig Alexa 488 | Jackson ImmunoResearch | 706-545-148 |
| Donkey anti-guinea pig Cy3 | Jackson ImmunoResearch | 706-165-148 |
| Donkey anti-guinea pig Alexa 647 | Jackson ImmunoResearch | 706-605-148 |
| Software and Algorithms |  |  |
| Prism 9 | GraphPad | N/A |
| Imaris | Bitplane, Switzerland | N/A |
| CellRanger | 10X Genomics | N/A |
| R Statistical Computing Software version 4 | R Project | N/A |
| RStudio | RStudio, Inc | N/A |
| Python 2.7 | Anaconda, Inc | N/A |
| Python 3.6 | Anaconda, Inc | N/A |

### Determination of candidate selector TFs

We used Seurat v3^62^ to determine the sets of differentially expressed TFs that are continuously maintained in each neuron from our developmental scRNA-seq atlas of the optic lobes^5^. First, we subset Seurat objects from each stage to retain only the neuronal clusters. Cluster markers were then calculated between these neuronal clusters only using the FindAllMarkers function (only.pos=TRUE, otherwise default parameters) which we filtered for transcription factors using a list of 629 genes from FlyBase. From these tables, we determined for each cluster, the TFs that were picked as its markers in adults as well as in at least 4 out of the 5 developmental stages (to buffer against potential errors due to technical variations in small clusters). A total of 113 TFs were found to be consistent markers of at least one cluster throughout development and in adults^5^.

Relying on cluster markers alone can be misleading: Some genes may fail to be picked up as markers for certain clusters despite being expressed if the level of expression is not significantly higher than the average of the rest of the dataset. This could be true even for differentially expressed genes. Similarly, some pan-neuronal (or ubiquitous) genes might be picked up as markers in some clusters if the level of expression is variable. We thus turned to mixture modeling data that we previously generated to binarize the expression of every gene in each cluster and stage^5^. This method considers, for each gene, the level of expression in all clusters in order to assign a probability of expression to each cluster^48^. For each of the 174 neuronal clusters with mixture modeling available at all 6 stages, we determined which of the 113 marker TFs had a non-zero probably of expression for a given cluster at every stage. However, this can also be error-prone for some genes when there is not a bimodal distribution of expression levels among the clusters. Therefore, we combined the two methods: a gene was selected as a candidate selector TF for a given cluster if i) it was found as a consistent marker of the cluster as described above, <u>or</u> ii) if it was continuously expressed at all stages according to mixture modeling. Lastly, we discarded all TFs that were found to be expressed in more than 150 clusters according to mixture modeling (likely pan-neuronal) in at least two of the stages.

The final table (**Fig. S1A**, **Table S1**) contains 95 TFs in total, with a median of 10 TFs per cluster (min:4, max:16) whose combinations are unique to each cluster.

It is important to note that while these results represent our best efforts to binarize a complex developmental gene expression landscape, there may still be errors, especially for genes with high variations in their levels of expression, and for smaller clusters that may be present in less than 20-30 cells in some of the stages. We thus recommend manual inspection of candidate selector expression patterns for cell types of interest as we did here (**Fig. 1B**, **Fig. 3D**) before designing experiments.

### Animal husbandry and genetics

All experiments were performed with both female and male (mixed) *D. melanogaster* (except for sequencing experiments, where only females were used) maintained at 25 or 29°C. The precise genotypes and temperatures used for experiments in each figure panel are detailed in **Table S3**. Generally, MARCM and sequencing experiments were performed at 25°C, while RNAi and most ectopic expression experiments were performed at 29°C. Source details for all fly strains are specified in the Key Resources Table. Adult dissections were performed within 2 days of eclosion. For pupal dissections, white pupae (P0) were selected and then aged until P50 (50h at 25°C or ∼35h at 29°C) or P30 (30h at 25°C or ∼20h at 29°C). Late L3 experiments were performed using wandering larvae. Experiments involving tub-Gal80^ts^ were reared at 18°C and then switched to 29°C two days before dissection (L3 dissections) or at late L3 stage (later dissections). Heat shocks at 37°C were applied for 15 minutes at L3 stage (MARCM) or for 7- 8 minutes at early pupal (P0-10) stages (flip-out).

### Antibody generation

Polyclonal antibodies against Drgx, Pdm3, SoxN, Mef2, Vsx2, Brp and Repo were generated by Genscript (https://www.genscript.com/).

Amino acids 431-587 of Drgx protein was used to immunize rats:

SNSVAELRRKAQEHSAALLQSLHAAAAAGLAFPGLHLPPLSFAHHPALGQHVVNHNNNNTMRMKHEAQDMTMNGLGPGSGSGSGSGSAGGGTSSAALLDLAESAVAYQQQQHATLSPPTTPTQQSSGGVAATEGSPGSGAIAGSGSLNGNVVLTKME

Amino acids 801-1292 of Pdm3 protein was used to immunize rats:

STSAVSSTLPQISLRHPDELTAPQMDLKPLELSASTSPPAPPPRHHFGHSLRGSSTVSPKHS PQGRMGGSGGSTTTGMNLSQHHERHDRLERLERQERHERRSHTPTATATRASVSSSSSA GHHGGSLPSGRLSPPSSAPSNSAANSISDRGYTSPLFRTHSPQGHALSLGGSPRLERDYLG NGPSSGTATSTSSCGAPTAAGSSATANVLSSINRLNASNGELTITKSLGAPTATATRASSASP RDDSPGPGPSTSSVSHMQPLKLSPSSRSEPPHLSPNGNDNDNDLLMDSPNEPTINQATTNV VDGIDLDEIKEFAKAFKLRRLSLGLTQTQVGQALSVTEGPAYSQSAICSSALAAQMYAAQLST QQQNMFEKLDITPKSAQKIKPVLERWMKEAEESHWNRYKSGQNHLTDYIGVEPSKKRKRRT SFTPQALELLNAHFERNTHPSGTEITGLAHQLGYEREVIRIWFCNKRQALKNTVRMMSKGMV

The following peptide from SoxN protein^82^ was used to immunize rabbits:

LHYQTDSPDLQQQHQS**C**

The entire ORF of the Mef2 protein purified with a His-tag was used to immunize rabbits:

MGRKKIQISRITDERNRQVTFNKRKFGVMKKAYELSVLCDCEIALIIFSSSNKLYQYASTDMDR VLLKYTEYNEPHESLTNKNIIEKENKNGVMSPDSPEAETDYTLTPRTEAKYNKIDEEFQNMMQ RNQMAIGGAGAPRQLPNSSYTLPVSVPVPGSYGDNLLQASPQMSHTNISPRPSSSETDSGG MSLIIYPSGSMLEMSNGYPHSHSPLVGSPSPGPSPGIAHHLSIKQQSPGSQNGRASNLRVVI PPTIAPIPPNMSAPDDVGYADQRQSQTSLNTPVVTLQTPIPALTSYSFGAQDFSSSGVMNSA DIMSLNTWHQGLVPHSSLSHLAVSNSTPPPATSPVSIKVKAEPQSPPRDLSASGHQQNSNG STGSGGSSSSTSSNASGGAGGGGAVSAANVITHLNNVSVLAGGPSGQGGGGGGGGSNGN VEQATNLSVLSHAQQHHLGMPNSRPSSTGHITPTPGAPSSDQDVRLAAVAVQQQQQQPHQ QQQLGDYDAPNHKRPRISGGWGTHHHHHH

The following epitope from Vsx2 protein was used to immunize rabbits:

TEAPTDLTTTAGATVAKERQTPTPPKTTNATMATAATSAATAATPTNAAEGNLTSVSEPQQQ PQQQQQEQQHHQPHHHQYREHHQMTMAAASRMAYFNAHAAVAAAFMPHQLAAAVHHHH QHQHQHHPHHHPHHPHGAVGGPPPPPPMQHHHPHHPHHPLLHAQGFPQLKSFAAGAGTC LPGSLAPKDFGMESLNGFGVGPNSKKKKK

Amino acids 221-401 of Brp protein was used to immunize guinea pigs:

TDVQRQQLEQQQKQLEEVRKQIDNQAKATEGERKIIDEQRKQIDAKRKDIEEKEKKMAEFDVQLRKRKEQMDQLEKSLQTQGGGAAAAGELNKKLMDTQRQLEACVKELQNTKEEHKKAATETERLLQLVQMSQEEQNAKEKTI MDLQQALKIAQAKVKQAQTQQQQQQDAGPAGFLKSFF

The following epitope from Repo protein was used to immunize guinea pigs:

MEHDSFDDPIFGEFGGGPLNPLGAKPLMPTTTAMHPVMLGSVHELCSQQQQQQQQQRLPD CNTILPNGGGGGAGSGGAGG SPNYVTKLDFVNKMGCYSPSQKYEYISAPQKLVEHHHHHH

### *Drgx*^ΔTm^^1^ allele

CRISPR-mediated mutagenesis was performed (WellGenetics Inc) to produce a genomic deletion of the Tm1-specific enhancer element we identified in the *Drgx* locus (see below the scATAC-seq analysis section). In brief, the upstream gRNA sequence AGCAATTCGCTATCCTTCGC[TGG] and downstream gRNA sequence TTCGACTGGACAGCTTAGTC[TGG] were cloned into U6 promoter plasmid(s) separately. The cassette 3xP3-RFP, which contains a floxed 3xP3-RFP, and two homology arms were cloned into pUC57-Kan as donor template for repair. gRNAs and hs-Cas9 were supplied in DNA plasmids, together with donor plasmid for microinjection into embryos of the control strain w^1118^. F1 flies carrying selection marker of 3xP3-RFP were further validated by genomic PCR and sequencing. CRISPR generated a 1,038 bp deletion within the 4^th^ intron of *Drgx*. 3xP3-RFP cassette was then excised through Cre/LoxP recombination, and the excision was validated by PCR and sequencing.

### Immunohistochemistry and imaging

Fly brains were dissected in ice-cold Schneider’s Insect Medium (SIM) and fixed in 3.7% formaldehyde (in PBS) at room temperature for 30-50 minutes and washed in PBST (PBS + 0.3% Triton X-100). They were then incubated in primary antibodies (in PBST + 5% horse serum) for 1 or 2 days at 4°C, washed three times in PBST for 10 minutes and then incubated in secondary antibodies at 4°C overnight followed by 2x 15 min washes in PBST and an additional 15 min in PBS. They were then mounted in Slowfade and imaged with a Leica SP8 confocal microscope using a 63x (NA=1.3) glycerol objective with xy resolution <200nm and z resolution = 500nm.

The following primary antibodies were used: Chicken anti-GFP (1:1000), mouse anti-V5 (1:250), rat anti-NCad (1:20), mouse anti-Brp (nc82, 1:50), guinea pig anti-Brp (1:300), mouse anti-Aop (1:100), rabbit anti-SoxN (1:250), rabbit anti-Mef2 (1:250), rat anti-Drgx (1:250), rat anti-Pdm3 (1:500), guinea pig anti-Pdm3, guinea pig anti-Repo (1:500), rabbit anti-Vmat (1:250), guinea pig anti-Vsx1 (1:300), rabbit anti-Vsx2 (1:500), guinea pig anti-Runt (1:500), rabbit cleaved anti-Dcp-1 (1:200), rabbit P-p44/42 anti-MAPK (1:100).

The following secondary antibodies (all donkey) were used: anti-chicken Alexa 488, anti-rabbit DyLight 405, anti-rabbit Alexa 488, anti-rabbit Cy3, anti-rabbit Alexa 647, anti-mouse Alexa 488, anti-mouse Alexa 555, anti-mouse Alexa 647, anti-rat Alexa 488, anti-rat Cy3, anti-rat Alexa 647, anti-guinea pig DyLight 405, anti-guinea pig Alexa 488, anti-guinea pig Cy3, anti-guinea pig Alexa 647. All were used at 1:300 dilution except for anti-chicken Alexa 488 that was used at 1:500.

The origin of all antibodies used is listed in the Key Resources Table.

### Image analysis

Images were analyzed and prepared using Imaris 7 (Bitplane) and Photoshop 2021 (Adobe). They were presented either as maximum intensity projections to highlight TF staining patterns or as 3D reconstructions using the “Blend” mode with “glow” colormap in Imaris to highlight neuronal morphologies.

Most quantifications were done manually by determining the proportion of each cell type (ignoring irrelevant cell types that may be labeled by the drivers, such as non-Tm neurons in experiments related to Tm1/2/4/6) in each brain (biological replicates) based on their stereotypical morphology, or expression of specific TFs, as specified in the figure legends. As the number of labeled neurons per brain show high variation in MARCM and flip-out experiments, all statistics were performed using the per brain proportions of indicated cell types (or their perturbed variations) and parametric, two-sided t-tests unless indicated otherwise. Bar and pie charts were generated using Prism 9 (GraphPad).

Quantification of the number of apoptotic cells for experiments shown in **Figure S2G** was performed semi-automatically using Imaris. Colocalization channels were created between the GFP (min. intensity=30) and Pdm3 (min. intensity=50) channels, as well as between the GFP (min. intensity=30) and cDcp1 (min. intensity=30) channels. Surface objects were then created separately for the i) GFP, ii) GFP-Pdm3 Coloc, iii) GFP-cDcp1 Coloc channels, with the same thresholds, filtered for “Quality” above 5 and “Number of Voxels” above 500. The number of cells that co-express GFP+Pdm3+cDcp1 were counted manually based on the overlap of Surface objects from (ii) and (iii). For each brain, the ratio of all dying cells were then calculated as the number of Surface objects in (iii) divided by the number of objects in (i). The ratio of dying Tm2 neurons were calculated as the ratio of triple labeled cells divided by the number of objects in (ii). The analysis was restricted to the medulla cortex as T2 and TE neurons are also labeled by GFP and Pdm3 in this experiment, but they were excluded based on their location.

### Single-cell RNA sequencing

Female white pupae (P0) were selected and maintained at 25°C until P50. Brains were dissected in ice-cold SIM and incubated for 20 minutes (at 25°C) in a Collagenase/Dispase cocktail, followed by washes and mechanical dissociation of the optic lobes, using a previously described protocol^5^. 5 brains (10 optic lobes) were dissociated together for each condition (Control and UAS-Pdm3). We then isolated the cells expressing the nuclear GFP *Stinger* from these single-cell suspensions using a BD FACSAria (see **Supplementary Information** for the FACS gating strategy). Samples were immediately processed for scRNA-seq.

Droplet-based purification, amplification and barcoding of single-cell transcriptomes were performed using Chromium™ Single Cell 3’ Reagent Kit v3.1 (10X Genomics) as described in the manufacturer’s manual. As the output concentrations from FACS were typically less than 400 cells/μl, we loaded the entire reaction volume (43.2 μl) with single-cell suspension (no extra water). The samples of each condition were processed back-to-back at each step of the entire protocol and were run on the same 10x chip to minimize batch effects. The libraries were subjected to paired-end sequencing (28x8x91) with Illumina NovaSeq 6000 (Genomics Core at NYU CGSB) to ∼130,000 reads per cell (88-89% sequencing saturation, as reported by CellRanger). We mapped the sequenced libraries to the *D. melanogaster* genome assembly BDGP6.88 using CellRanger 5.0.1, which retained 3,830 cells from Control and 3,813 cells from UAS-Pdm3 library, with a median of ∼2500 genes per cell. All downstream analyses were performed with Seurat v4^83^. The libraries were merged (without integration) and further filtered to retain the cells with 2,000 to 20,000 UMIs and less than 5% of UMIs corresponding to mitochondrial genes, to a total of 6693 cells (3303 Control, 3390 UAS-Pdm3).

### scRNA-seq analysis

The data was log-normalized, followed by determination of the top 2000 variable features and scaling with default parameters in Seurat. Cell type assignments were done in a supervised manner using a machine learning model we had previously trained on the scRNA-seq atlas of the entire optic lobe at P50^5^. tSNE reduction was then calculated using the top 50 principal components (**Fig. S2F**). Surprisingly, these libraries contained many cell types which we never observed using the *TmX/Tm2-Gal4* with a membrane-bound GFP (**Fig. S2D**), including (but not limited to) Mi1/4/9 and Tm3/5/9/20 types (**Fig. S2F**). It is possible that the driver has extremely weak background expression in these other cell types, and that the very strong nuclear localization of *Stinger* allows GFP signal to be concentrated there (as opposed to a membrane GFP) allowing these cells to be picked up by FACS. Indeed, a high number of nuclei with weak GFP signal (that were never labeled by Pdm3 or Aop) could be observed in these brains (**Fig. S2G**). We thus restricted all further analysis to the cells classified as Tm1/2/4/6 or T2 neurons by the machine learning model (**Fig. 1H,J**). In addition, we realized that one of the brains dissected for the Control library was in fact a male, based on *roX1/2* expression. These cells were excluded from all the differential gene expression analyses below.

They were, however, still included in the UMAPs since those were calculated using the scaled data slot, which was corrected (regressed out) based on *roX1* and *roX2* expression.

The top 500 variable features were re-determined after subsetting of the data, which were then re-scaled accordingly. The UMAP reduction (**Fig. S2H**, **Fig. 1H-O**) was calculated using the top 6 principal components. The unsupervised clustering was performed using the top 5 principal components and resolution 0.4 (otherwise default parameters). Differentially expressed genes (DEGs) between the indicated clusters in **Figure 1K** and **Table S2** were then determined using the FindMarkers function with default parameters, filtered to only retain the markers with adjusted p-value < 0.01. Control and UAS-Pdm3 libraries in this experiment are expected to have some batch effects. To avoid picking up DEGs which may be driven by such effects, we also calculated the list of markers (adj. p-value < 0.01) between the 4 Tm types in the *control library only*, using the FindAllMarkers function (only.pos=TRUE, otherwise default parameters). Reasoning that any gene that does not normally display differential expression between the Tm neurons should not be regulated by Pdm3 in these cell types, the number of DEGs displayed in **Figure 1K** and the full marker lists provided in **Table S2** only include such genes that were also part of this list of 1395 “Control Tm markers”.

### scATAC-seq analysis

We used Seurat v4 and Signac 1.3.0^84^ to analyze the scATAC-seq datasets generated from whole fly brains by^27^ at stages Adult, P48 and P24. Peak/count matrices generated by the authors, based on pre-determined regulatory regions in the fly genome, were extracted from the CisTopic object and separate Seurat/Signac objects were created for all three stages. The following steps were then taken separately for each object/stage: The data were TF-IDF normalized and all features were set as variable by running the FindTopFeatures function with “min.cutoff = ‘q0’” option. Latent semantic indexing (LSI) reduction was then calculated using the RunSVD function (n=200). We then clustered the data using the top 120 LSI dimensions (excluding the first one, which correlated strongly with sequencing depth) and with ‘algorithm’ and ‘resolution’ set to 3 in the FindClusters function. Label transfer was applied from Adult to P48 and from P48 to P24 datasets to establish (rough) linkages between the clusters of different stages. The anchors were found between the datasets with 100 CCA dimensions, then the labels were transferred with the TransferData function (weight.reduction=lsi and dims=2:80).

In order to annotate the datasets, gene activity, i.e. ‘pseudo expression’, matrices were calculated by counting the number of reads (directly from the fragment files) within 5kb upstream of TSS for every gene. These were then used to perform label transfer from the scRNA-seq datasets at corresponding stages: Adult, P50 and P30^5^. The anchors were found between the datasets using the variable features of the RNA dataset (CCA, 150 dims), then the labels were transferred with the TransferData function (weight.reduction=lsi and dims=1:120). The quality of label transfer was variable and particularly low at P24; we thus first focused on the Adult dataset to establish correspondence between the scRNA- and scATAC-seq clusters. Every cluster was inspected manually for its correspondence to the optic lobe clusters in the scRNA-seq dataset. If a substantial number (depending on the expected frequency of clusters) of cells belonging to any (neuronal) RNA cluster corresponded to a clearly defined ATAC cluster, those clusters were retained for further analysis (**Fig. S4A**). This process excludes all glia and the vast majority of central brain neurons, but it may also exclude optic lobe neurons that derive from central brain which are typically present in lower frequencies and may not cluster well. Using the linkages we established above between the different stages, we determined in P48 and P24 datasets the clusters that corresponded to optic lobe neuronal clusters in the Adult dataset and discarded the rest.

After isolating the optic lobe neurons in all 3 stages, we reapplied TF-IDF and SVD, and re-clustered the datasets to maximize resolution. The Adult optic lobe dataset was clustered with LSI dimensions 3 to 80 and resolution=5, while we used the dimensions 2 to 80 and resolution=3 for the P48 and P24 datasets. Multiple clusters corresponding to T4/T5 neurons were merged. We obtained 45 final clusters of optic lobe neurons at the adult stage, 37 of which could be confidently linked to defined (though not all annotated) clusters (or groups of clusters) in the scRNA-seq atlas (**Fig. S4B**, the clusters that are still likely optic lobe neurons but could not be confidently linked to well-defined RNA clusters are grayed out). There were 59 initial clusters in the P48 dataset and 41 in P24. We took a multi-step process to harmonize the cell type assignments between different stages. First, label transfer was performed from Adult to P48 dataset, as described above for the whole datasets. We took advantage of both these transferred labels and the transferred labels from the P50 RNA dataset to manually inspect all clusters. Unlike the strategy we previously employed to harmonize the cell-type assignments in scRNA-seq datasets of different stages^5^, label transfer between different stages was far less robust in scATAC-seq; thus we mostly respected the unsupervised assignments at each stage. If most (typically >50% but in a few cases less) of a P48 cluster was classified as a specific Adult cluster (and no other), the entire cluster was assigned the same number as in the Adult. Since the clustering resolution was higher at P48, that meant in several cases merging of clusters. However, we later subclustered the lamina neurons L1-5 at this stage, to resolve all 5 types (**Fig. S4C**). In some cases where a P48 cluster was mapped to multiple Adult clusters, the P48 cluster was split based on the label transfer classifications. In other cases where the correspondence of a P48 cluster to the Adult clusters was unclear, they were assigned to new identities (not present in the Adult dataset). This was the case for TE neurons (that are not present in adult brains), but also for TmY14, Lawf and Lai neurons, which could not be mapped reliably in the Adult dataset. The remaining P48 clusters that had inconsistent mapping to Adult clusters and also could not be reliably linked to a P50 RNA cluster were kept but grayed out in tSNE (**Fig. S4C**). The process was then repeated from P48 to P24 dataset, which had the lowest clustering resolution as well as the least reliable mapping from the (P30) RNA dataset. This meant that most of the clusters had to be split manually based on the transferred labels from the P48 dataset to maintain the same resolution (**Fig. S4D**). For the same reason, a much larger proportion of the dataset remained unlinked to specific cell types and were grayed out in the tSNE.

The accessibility tracks were generated for the indicated clusters (**Fig. 2D**, **Fig. S4E**) using the CoveragePlot function with default parameters. The sequence of the genomic range chr2L:3679100-3679800 which encompasses the Tm1-specific peak in *Drgx* locus was extracted from Ensembl for motif enrichment analysis. We compiled a list of 1288 TF binding motifs, representing 411 TFs, from the CisBP database, including only those determined using direct evidence from *Drosophila*. We used the MEME suite (https://meme-suite.org/meme/tools/ame) tool AME^85^ to identify the enriched motifs. The only motifs that were found to be enriched with an E-value < 100 belonged to the TFs *Klu*, *org-1* and *nau*; the latter two are not expressed in the optic lobe at any stage according to our scRNA-seq datasets.

### Network Inference

We inferred GRN models for two distinct groups of neurons using data from developmental stages P24-P50: i) Tm1/2/4/6 neurons that we studied in this paper (Tm network), ii) Lamina monopolar neurons L1-5 whose development have been extensively studied previously (Lamina network).

#### Construction of priors

Prior connectivity matrices (tables of 1s and 0s linking TFs to target genes) were determined separately for Tm and Lamina neurons using the P48 scATAC-seq data (**Fig. S4C**), Signac v1.3.0 and the Inferelator-Prior software^58^ available on GitHub, from the release branch v0.2.3 faf5e47.

For each prior, we used GTF and fasta files from the *D. melanogaster* genome assembly BDGP6.88. MEME suite tool FIMO^86^ was implemented with the default 1e-4 p-value threshold to scan the *Drosophila* genome for the set of TF motifs described in the previous section. Several TFs motifs, specifically those belonging to: ‘ct’, ‘dan’, ‘danr’, ‘Eip78C’, ‘H2.0’, ‘jumu’, ‘Sox100B’, ‘Tbp’, ‘vis’, ‘Met’, were never found in the genome using the default FIMO threshold (likely due to their lower information content), thus for those motifs we used a 1e-3 p-value threshold to receive scores. All TF motifs were scored and filtered based on their information content as part of the Inferelator-Prior workflow. The refined list of scored TF motifs was used to assign regulatory relationships to target genes by using a 10kb window up and downstream of each gene body. These interactions were then clustered using the sklearn DBSCAN package with parameter epsilon=1 to retain a very sparse (<0.5% density on average per TF) matrix containing only the highest confidence interactions. The command line arguments for the Inferelator-Prior pipeline can be accessed at: https://github.com/cskokgibbs/DMOLN_NetworkScripts. This pipeline requires as input a .bed file defining genomic ranges to be scanned for motifs. We tested 3 different priors (NoBed, ChromA and MergedDA) for both Tm and Lamina networks whose underlying peak sets were determined with different strategies as described below. Please note that even though **Figure S6** displays the benchmarking results for networks built with all 3 priors, the final networks used in this work were based on the MergedDA priors.

##### NoBed

No constraining peak file was used to build these priors. The entire 10kb up/down window for every gene was scanned for motifs. Note that this implies the NoBed priors for Tm and Lamina networks were therefore identical.

##### ChromA

For these priors, we extracted all fragments originating from the Tm or Lamina neurons in the P48 scATAC-seq (**Fig. S4C**, red and blue ellipses) and called peaks using ChromA^87^ version: fcfd120 branch: Any-Genome, using the parameters: atac -spec dm6. These peak sets were then used directly as input to Inferelator-Prior. Therefore, for these priors, all regions accessible in any of the 4 Tm or 5 lamina neurons were scanned for motifs without considering differential accessibility.

##### MergedDA

We determined differentially accessible (DA) regions between the Tm or Lamina neurons with two different approaches. First, we used directly the peak set from^27^. We subset the P48 Signac object for either the 4 Tm clusters or the 5 lamina clusters (**Fig. S4C**) and calculated the DA peaks using the FindAllMarkers function (logfc.threshold = 0.1, test.use = ‘LR’, latent.vars = ‘nCount_peaks’). However, we noticed that these pre-determined regulatory regions often contained multiple peaks whose accessibility can vary independently, which could introduce artifacts. With this in mind, we re-built the peak/count matrices for Tm and Lamina clusters from the Fragments file using the FeatureMatrix function, using as features the peaks determined above by ChromA as accessible in Tm or Lamina clusters. DA peaks were then calculated for each group using the same parameters. This approach may have its own limitations, primarily due to sensitivity of peak calling since this was done separately on relatively small groups of cells here rather than the entire dataset. We thus decided to merge the two peak sets (separately for Tm and Lamina) using the GenomicRanges function reduce (min.gapwidth = 0) to obtain the final as input to Inferalator-Prior.

MergedDA networks outperformed the other priors in all metrics (**Fig. 5B**, **Fig. S6A-B**), as expected; however, ChromA and NoBed priors performed similarly, suggesting that ATAC-seq datasets are only useful for network inference when considering differential accessibility.

#### Inference

We used the scRNA-seq datasets from P30, P40 and P50 generated by our lab^5^, as well as from P24, P36 and P48 generated by another group^61^ to infer GRN models. In order to determine the sets of differentially expressed genes (DEGs) to model on, we used exclusively the “DGRP” portion of the P24-P36-P48 datasets as these are not expected to have batch effects between stages. The objects were subset for either the Tm or Lamina clusters, and then the stages were merged to keep each individual cluster from each stage as a separate class (thus, 12 classes for Tm, 15 for Lamina). DE genes (adj. p-value<0.01) were then calculated between these classes using the FindAllMarkers function (only.pos = TRUE). The resulting sets of DE genes therefore represent those whose expression vary either across different cell types and/or different stages. Log-normalized data was used for inference; however, applying the same default scaling factor (10,000) to all stages can introduce artifacts if there are variations in the total RNA content of a given cell across different stages of development. We previously noted significant differences in the average UMI per cell values across different stages of our dataset^5^; though it remains unclear how much of this could be attributed to technical reasons. When we normalized the cells of each stage with a value proportional to the average UMI value of that stage, we received overall very similar results compared to when we normalized with the default value. However, the benchmarks we performed with the *Hr3* RNAi dataset (described below) were significantly better (results not shown) with the stage-adjusted normalization; we thus proceeded with this method. This is consistent with the finding that *Hr3* is strongly regulated in the temporal axis during this period^14^.

The networks were inferred using the Inferelator 3.0^58^, GitHub version v0.5.6 dd532f4, branch: release. Networks were learned using the multitask framework “AMuSR”^88^ with the following parameters: regression=”amusr”, workflow=”multitask”, bootstraps=5, and cross validation seed set to 42 . Datasets were organized into two tasks representing data collected at P30-40-50 (“Desplan”) and P24-36-48 (“Zipursky”). For each task, TFA was calculated by computing the dot product between the task-specific expression dataset and the pseudo- inverse of the prior. As part of the inference step, a linear model for each task is constructed that assumes each gene in the expression dataset can be modeled as a linear combination of the TFs regulating it (**X** = **βA**). Task specific betas represent the variance explained by each regulatory TF and are solved in the following way: We calculate the ratio between the variance of the residuals in the full model and the variance of the residuals when the model is refit, leaving one TF out at a time, as originally explained in the Inferelator^58^. Any non-zero beta is considered evidence for a regulatory relationship between a gene and a TF. In the multi-task learning approach, after TFA is computed for each task independently, we solve the linear model for each task together resulting in an ensemble network, as well as task specific networks. Here, the model implements adaptive penalties to prefer regulatory interactions shared across the two tasks, placing a heavier penalty on interactions found to be only task-specific. Three networks were learned using the respective priors described above for both groups of cell types. In order to obtain results without sampling error, model selection was repeated by sampling the expression data across 5 bootstraps, resampled with replacement. Tm and lamina networks were modeled on both DE genes and DE TFs, as described above. The respective priors, subset by DE genes and TFs, were used as gold standard networks for evaluating network performance. Evaluation metrics included calculating Area Under the Precision Recall (AUPR) curve to compare our predicted interactions to the gold standard network (prior). Further, we compute Matthews correlation coefficient (MCC) to obtain confusion matrices representing true positives (TP), false positives (FP), true negatives (TN), and false negatives (FN). Calculating this score provided us with a metric to evaluate the reliability of our predicted network, as MCC obtains a high score only if the predicted interactions obtain good results for all four confusion matrices. An F1 score was additionally computed by taking the weighted average of the precision and recall. This score allows us to take into account both FP and FN, which is important to consider in the case of imbalanced classes, such as our model’s assumption that a target gene can be regulated by a limited number of TFs. Finally, we compute the variance explained by each edge contained in the resulting network, allowing us to interpret which edges explain the most variance in the posterior network. Each network was benchmarked against a negative control where a network was inferred on each respective prior with the gene names shuffled. The final (multitask, MergedDA prior) Tm and Lamina networks are provided in **Table S4**. All network inference run scripts for the Inferelator are available at https://github.com/cskokgibbs/DMOLN_NetworkScripts.

The Inferelator does not report the sign (activating or repressing) of inferred interactions. Thus, for visualization purposes, we calculated Pearson correlations of expression levels using the stage-adjusted, log-normalized data with all the stages merged for the respective cell types. The network visualizations were generated with the Python package jp_gene_viz version 2.30.1 on branch ‘Master’ using the filters described in figure legends.

#### Benchmarking the networks with TF perturbations

As the Inferelator attempts to model all biologically relevant (between cell types and stages) differential gene expression among the input cells, this provides an opportunity to test if the learned models are generalizable to novel regulatory states never seen by the algorithm. We took advantage of 3 independent RNA-seq datasets acquired in the background of specific TF perturbations (described below) for this purpose. For each dataset, TFA was calculated for each cell using the respective priors (Lamina for *Hr3* and *Erm*, Tm for *pdm3* datasets) and the log-normalized expression values. We incorporated the betas collected from the relevant networks outlined above, describing the relative contribution of TFs to the model of each gene in our sample, to provide this benchmark with unseen, independent data. Taking the dot product of the estimated TFA and independent betas for each experiment provided us with reconstructed (predicted) expression matrices. The command line arguments for these steps are provided in the same Github repository referenced above. Note that since the models were built on DEGs only, the predicted expression matrices also contain only these genes (1394 genes for Tm, 1684 for lamina datasets). We evaluated the results produced by all three prior generation methods described above for these benchmarks (**Fig. S6H-J**) but the UMAPs shown are only for the reconstructions generated with the “MergedDA” priors (**Fig. 5D-F**, **Fig. S6E-G**).

##### Hr3 RNAi

The Seurat object provided by the authors^14^ was converted to match the gene names to our reference annotation, and subset to retain only the P48 stage. The predicted expression matrices were generated using the Lamina priors to calculate TFA, and the betas generated specifically from the “Zipursky” task described above, since these datasets contain the matching time point (P48) and were produced by the same lab using the same sequencing technology (10x v3.1). A merged Seurat object was then created with both the real (including only the DEGs present in the predicted expression) and predicted expression matrices, identification of top 1000 variable features, scaling and PCA were performed with the default parameters. UMAP visualization (**Fig. S6E**) was created using 30 principal components. The object was then split based on real vs. predicted expression, and integration was performed using the FindIntegrationAnchors (reduction=”cca”, anchor.features=1000, dims=1:30) and the IntegrateData (dims=1:30) functions. The data were scaled again and UMAP was generated using the same parameters as above (**Fig. 5D**). Then, separately for each of the 5 cell types contained in this dataset (L1-5), we determined the DEGs (adj. p-value<0.01) between control (w RNAi) and Hr3 RNAi conditions using both the real and predicted clusters (non-integrated data). For each cell type, precision is defined as the ratio of correctly determined DEGs between the conditions in predicted clusters (i.e., those that were also DEGs between the real clusters) (TP/(TP+FP)), and the recall is defined as the ratio of these correctly predicted DEGs to all DEGs between the real clusters (TP/(TP+FN)) (**Fig. S6H**).

##### Erm mutant

We obtained the Transcripts Per Million (TPM) results from a bulk RNA sequencing experiment of FACSed L3 neurons mutant for Erm (dFezf1) at stage P40^32^ and converted the gene names to match our reference annotation. We then simulated single-cell transcriptomes, separately for each of the 5 replicates of control (FRT40) and mutant experiments (**Fig. S6D**). The expression values were re-normalized to a total of 1 by dividing each value with the sum of all values for that sample. A probability distribution table was then determined based on cumulative sums: starting from 0 and increasing gradually to 1 for the last (8905^th^) gene in the dataset, with the amount of increase for each gene from the previous one being proportional to its level of expression. Thus, for all expressed genes, their value is higher than the previous gene in the list, while the value remains constant (from the previous one) if the gene is not expressed. For each simulated single-cell, the number of UMIs it will contain was randomly chosen from a normal distribution with mean=2500, s.d.=2000 and a minimum of 1000 UMIs. Each UMI was then assigned to a gene by picking a random number between 0 and 1, and determining the first occurrence of a number smaller than that in the probability distribution. Lastly, 100 simulated cells with at least 500 genes expressed were selected per replicate. Overall, the process aims to recapitulate the noisy sampling of mRNAs with high drop-out rates as characteristic of scRNA-seq experiments. Log-normalization, variable feature selection and scaling was applied using Seurat and the UMAP visualization (**Fig. S6D**) was created using the top 3 principal components. As this data contains only a single cell-type (with and without perturbation of only one TF), we observed that 3 PCs were enough capture all biological variation, as the inclusion of more PCs only resulted in increasing distances between the replicate clusters of the same condition.

The predicted expression matrices were then generated using the simulated single-cells, as described above for Hr3, but the betas were used from the “Desplan” task in this case since this task contains cells sequenced at the same stage (P40) as the Erm dataset. Integration between the real and predicted cells was performed using 200 anchors and 10 reduced dimensions (otherwise same as above) and the UMAPs were again calculated using the top 3 PCs (**Fig. 5E**, **Fig. S6F**). As there is only one cell type in this case, a single precision and recall value was determined for each tested prior (**Fig. S6I**). Interestingly, when we used the non-specific “NoBed” prior instead of the MergedDA prior, the predicted clusters in both Hr3 and Erm reconstructions tended to display many more DEGs between the conditions (higher recall), but with lower precision (**Fig. S6H-I**).

##### UAS-Pdm3

As both benchmarks above were on lamina neurons, we also aimed to similarly benchmark the Tm networks using the scRNA-seq dataset generated in this paper (**Fig. 1J**). TFAs calculated using the Tm prior, as well as the betas from the wild-type Tm networks (Zipursky task) were used to generate the predicted expression matrices. Normalization, scaling, dimensionality reduction and integration was performed using the same parameters as described above for the Hr3 dataset (**Fig. 5F**, **Fig. S6G**). Precision and recall metrics were calculated for the DEGs between indicated subclusters in **Figure S6J** (referring to the same cluster numbers as in **Fig. 1K**).

Relatively higher metrics for these reconstructions are likely a result of changes in this dataset representing complete transformations of one cell type into another, unlike the lamina datasets where “unnatural” regulatory states were created. The one exception to this was Tm1 subcluster 6, where DEG analysis against the control Tm1 cells revealed extremely low recall (**Fig. S6J**). As we have discussed in the first section, this cluster consists of cells that express ectopic *pdm3* (**Fig. 1N**), but still retain Tm1 fate and appear morphologically normal (**Fig. S2I**). It is therefore likely that the changes observed in these cells do not represent a significant shift in their regulatory state, potentially explaining the low performance of the model in reconstructing them.

#### GO Enrichment

We filtered the inferred GRN model of Tm neurons to only include the interactions with combined confidence greater than 80% and variance explained greater than 1% (the same filtering as in Fig. 5C). Within this filtered network, we determined the predicted targets of Hr3 (167 genes), Pdm3 (120 genes), Mef2 (112 genes) and subjected them (separately) to GO enrichment analysis for ‘Molecular Function’ terms using The Gene Ontology Resource (http://geneontology.org/). We filtered the terms to include only those with fold enrichment greater than 2. We used REVIGO (http://revigo.irb.hr/) to remove redundant terms and group the related ones^89^ with a similarity index of 0.5. The TreeMap tables were then exported from REVIGO and inputted to the Python package CirGO^90^ to create the summary graphs in **Figure S7E**.

