## Supplementary material for "Coordinated control of neuronal differentiation and wiring by a sustained code of transcription factors": FACS Gating Strategy

#### Supplementary Information: FACS Gating Strategy

BD FACSDiva 8.0.2

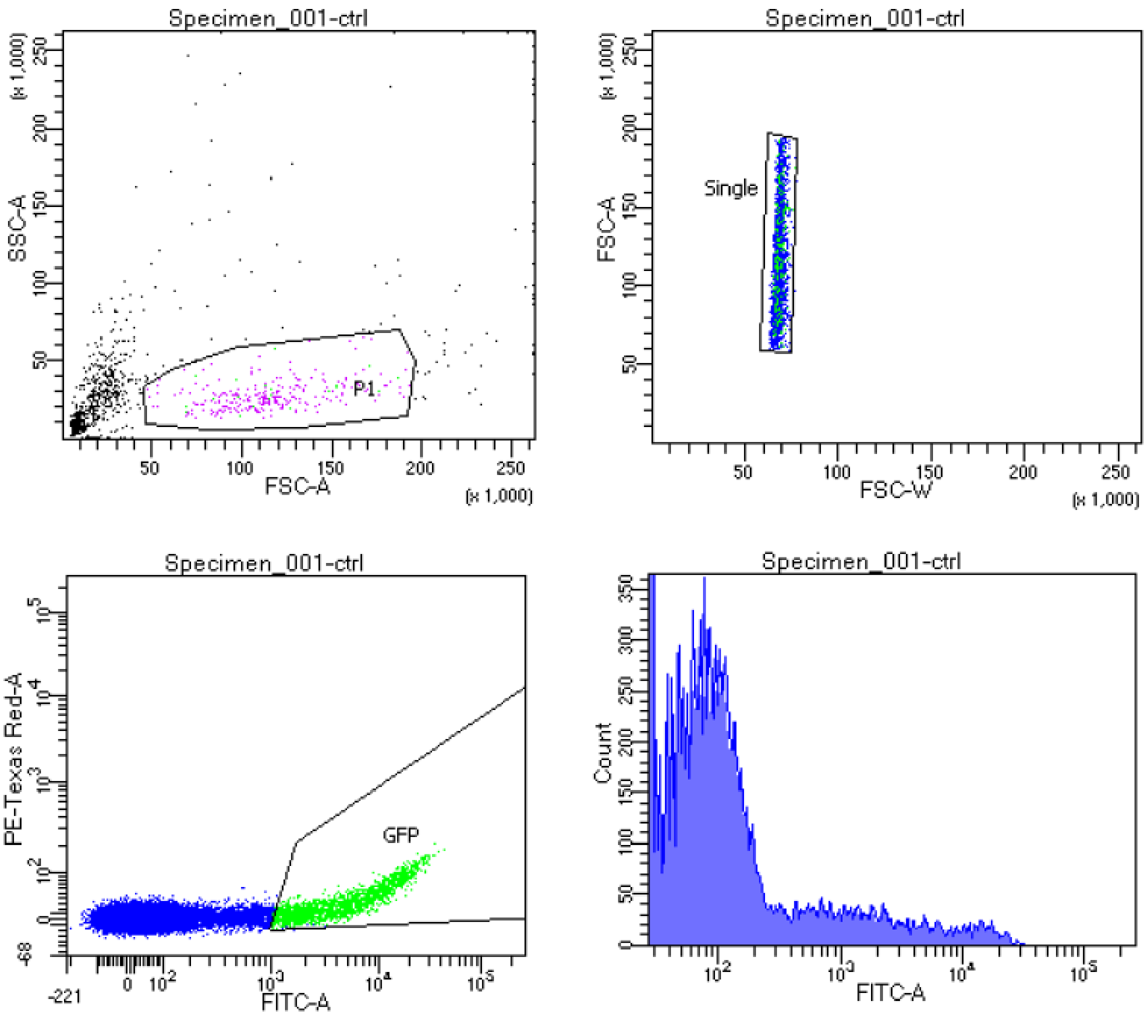


| | |
|---|---|
| Experiment Name: | Neset_006 |
| Specimen Name: | Specimen_001 |
| Tube Name: | ctrl |
| Record Date: | Sep 21, 2021 2:41:41 PM |
| $OP: | Administrator |
| GUID: | f6614b59-4cf6-4096-9a90-8c92... |

| Population | #Events | %Parent | DsRed-A Mean | DsRed-A Mean |
|---|---|---|---|---|
| All Events | 100,000 | #### | #### | 4 |
| P1 | 27,843 | 27.8 | #### | 2 |
| Single | 26,749 | 96.1 | #### | 2 |
| GFP | 2,059 | 7.7 | #### | 2 |

Tube: ctrl

| Population | #Events | %Parent | %Total |
|---|---|---|---|
| All Events | 100,000 | #### | 100.0 |
| P1 | 27,843 | 27.8 | 27.8 |
| Single | 26,749 | 96.1 | 26.7 |
| GFP | 2,059 | 7.7 | 2.1 |

### BD FACSDiva 8.0.2

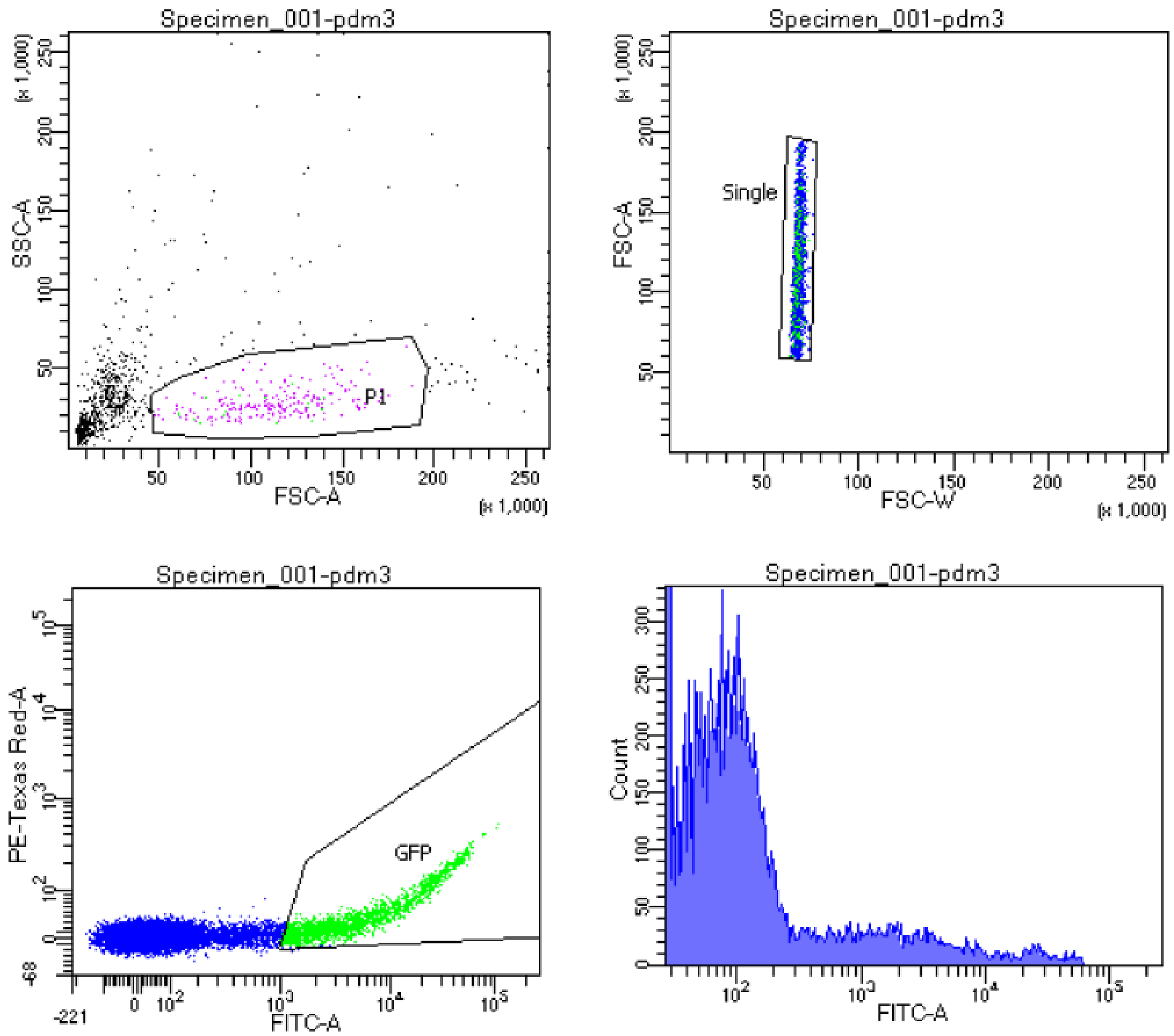


| | |
|---|---|
| Experiment Name: | Neset_006 |
| Specimen Name: | Specimen_001 |
| Tube Name: | pdm3 |
| Record Date: | Sep 21, 2021 2:53:50 PM |
| $OP: | Administrator |
| GUID: | 39f6ec45-bf38-40fd-ada3-8a3c... |

| Population | #Events | %Parent | DsRed-A Mean | DsRed-A Mean |
|---|---|---|---|---|
| All Events | 100,000 | #### | #### | 4 |
| P1 | 24,382 | 24.4 | #### | 3 |
| Single | 23,338 | 95.7 | #### | 3 |
| GFP | 2,054 | 8.8 | #### | 4 |

Tube: pdm3

| Population | #Events | %Parent | %Total |
|---|---|---|---|
| All Events | 100,000 | #### | 100.0 |
| P1 | 24,382 | 24.4 | 24.4 |
| Single | 23,338 | 95.7 | 23.3 |
| GFP | 2,054 | 8.8 | 2.1 |
