## Supplementary material for "Coordinated control of neuronal differentiation and wiring by a sustained code of transcription factors": Table S3

**Supplementary Table 3**

Ctrl: Control, Exp: Experiment, FSF: FRT-stop-FRT

| **Panels** | **Genotype** | **Temp** |
| --- | --- | --- |
| S2B, S3A-B, S5G | hsFLP2:PEST/+; UAS-FSF-CD4tdGFP/+; R35H01-Gal4/+ | 25 |
| S2C | hsFLP2:PEST/+; TI{CRIMIC.TG4.2}^Wnt10CR01661^/+ ; UAS-FSF-CD4tdGFP/+ | 25 |
| S2D, S5H | hsFLP2:PEST/+; UAS-FSF-CD4tdGFP/+; R71F05-Gal4/+ | 25 |
| 1C, S2E | Ctrl: hsFLP122, UAS-CD8GFP/+; FRT42D, tub-Gal80/FRT42D;  R71F05-Gal4/UAS-CD4tdGFP  Exp: hsFLP122, UAS-CD8GFP/+; FRT42D, tub-Gal80/FRT42D, pdm3^1^;  R71F05-Gal4/UAS-CD4tdGFP | 25 |
| 1D-F | Ctrl: hsFLP2:PEST/+; UAS-FSF-CD4tdGFP/+; R71F05-Gal4/+  Exp: hsFLP2:PEST/+; UAS-FSF-CD4tdGFP/ P{TRiP.HMJ21205};  R71F05-Gal4/+ | 29 |
| 1G | Ctrl: Same as above  Exp: hsFLP2:PEST/+; UAS-FSF-CD4tdGFP/ UAS-pdm3.short; R71F05-Gal4/+ | 25 |
| 1H-O, S2F-H | Ctrl: ;UAS-Stinger/+; R71F05-Gal4/+  Exp: ;UAS-Stinger/UAS-pdm3.short; R71F05-Gal4/+ | 25 |
| S2I | Ctrl: hsFLP2:PEST/+; UAS-FSF-CD4tdGFP/+; 27b-Gal4/+  Exp: hsFLP2:PEST/+; UAS-FSF-CD4tdGFP/UAS-pdm3.short; 27b-Gal4/+ | 25 |
| 2A-C,  S3D | Ctrl: hsFLP2:PEST/+; UAS-FSF-CD4tdGFP/+; R35H01-Gal4/+  Exp: hsFLP2:PEST/+; UAS-FSF-CD4tdGFP/ Mi{Hto-WP}Drgx^GLO^;  R35H01-Gal4/+ | 29 |
| S3C | hsFLP2:PEST/+; tub-Gal80^ts^/UAS-pdm3.short;  R35H01-Gal4/UAS-FSF-CD4tdGFP | 25 |
| 2E | ; tub-Gal80^ts^/+; pxb-Gal4, UAS-CD8GFP/+ | 29 |
| 2F | ; tub-Gal80^ts^/+; pxb-Gal4, UAS-CD8GFP/UAS-Klu.HA | 29 |
| 2G-I, S3E-F | Ctrl: hsFLP2:PEST/+; Drgx^ΔTm1^/+ ; 27b-Gal4/UAS-FSF-CD4tdGFP  Exp: hsFLP2:PEST/+; Drgx^ΔTm1^/Drgx^ΔTm1^; 27b-Gal4/UAS-FSF-CD4tdGFP | 25 |
| S3G | Ctrl: w^1118^ ;;  Exp: w^1118^ ; Drgx^ΔTm1^/Drgx^ΔTm1^ ; | 25 |
| 3A-B | Ctrl: hsFLP122, UAS-CD8GFP/+; FRT40A, tub-Gal80/FRT40A;  R35H01-Gal4/UAS-CD4tdGFP  Exp: hsFLP122, UAS-CD8GFP/+; FRT40A, tub-Gal80/FRT40A, SoxN^NC14^;  R35H01-Gal4/UAS-CD4tdGFP | 25 |
| S5A | hsFLP2:PEST/+; tub-Gal80^ts^/UAS-SoxN.V5;  R35H01-Gal4/UAS-FSF-CD4tdGFP | 29 |
| S5B | Ctrl: hsFLP122, UAS-CD8GFP/+; FRT40A, tub-Gal80/FRT40A;  R35H01-Gal4/UAS-CD4tdGFP  Exp: hsFLP122, UAS-CD8GFP/+; FRT40A, tub-Gal80/FRT40A, aop^E833^;  R35H01-Gal4/UAS-CD4tdGFP | 25 |
| S5C | Ctrl: hsFLP2:PEST/+; UAS-FSF-CD4tdGFP/+; R35H01-Gal4/+  Exp: hsFLP2:PEST/+; UAS-FSF-CD4tdGFP/+;  R35H01-Gal4/ P{TRiP.HMS01256} | 29 |
| 3C | Ctrl: hsFLP2:PEST/+; UAS-FSF-CD4tdGFP/+; R71F01-Gal4/+  Exp: hsFLP2:PEST/+; UAS-FSF-CD4tdGFP/+;  R71F01-Gal4/ P{TRiP.HMS01256} | 29 |
| 3F-G, S5E | Ctrl: hsFLP2:PEST/+; UAS-FSF-CD4tdGFP/+; R76F01-Gal4/+  Exp: hsFLP2:PEST/+; UAS-FSF-CD4tdGFP/UAS-Vsx1; R76F01-Gal4/+ | 29 |
| S5D | hsFLP2:PEST/+; UAS-FSF-CD4tdGFP/UAS-Vsx2.CC; R76F01-Gal4/+ | 29 |
| 4A,E | Ctrl: hsFLP2:PEST/+; UAS-FSF-CD4tdGFP/+; R71F05-Gal4/+  Exp: hsFLP2:PEST/+; UAS-FSF-CD4tdGFP/P{TRiP.HMS01691};  R71F05-Gal4/+ | 29 |
| 4B,F | hsFLP2:PEST/+; UAS-FSF-CD4tdGFP/UAS-aop.WT; R71F05-Gal4/+ | 29 |
| 4C,G | hsFLP2:PEST/+; UAS-FSF-CD4tdGFP/UAS-aop.ACT; R71F05-Gal4/+ | 29 |
| 5H, S7A | Ctrl: hsFLP122, UAS-CD8GFP/+; FRT42D, tub-Gal80/FRT42D;  R71F05-Gal4/UAS-CD4tdGFP  Exp: hsFLP122, UAS-CD8GFP/+; FRT42D, tub-Gal80/FRT42D, Hr3^K10308^;  R71F05-Gal4/UAS-CD4tdGFP | 25 |
| 5J | Ctrl: hsFLP2:PEST/+; UAS-FSF-CD4tdGFP/+; R71F05-Gal4/+  Exp: hsFLP2:PEST/+; UAS-FSF-CD4tdGFP/tub-Gal80^ts^;  R71F05-Gal4/UAS-Blimp1 | 29 |
| 5K | Ctrl: hsFLP2:PEST/+; UAS-FSF-CD4tdGFP/+; 27b-Gal4/+  Exp: hsFLP2:PEST/+; UAS-FSF-CD4tdGFP/+; 27b-Gal4/P{TRiP.HMS00924} | 29 |
| S7C | Ctrl: hsFLP2:PEST/+; UAS-FSF-CD4tdGFP/+; R71F05-Gal4/+  Exp: hsFLP2:PEST/Mi{Hto-WP}ct^BRO^; UAS-FSF-CD4tdGFP/+; R71F05-Gal4/+ | 29 |
